# Informing shigellosis prevention and control through pathogen genomics

**DOI:** 10.1101/2021.06.09.447709

**Authors:** Rebecca J. Bengtsson, Adam J. Simpkin, Caisey V. Pulford, Ross Low, David A. Rasko, Daniel J. Rigden, Neil Hall, Eileen M. Barry, Sharon M. Tennant, Kate S. Baker

## Abstract

*Shigella* spp. are the leading bacterial cause of severe childhood diarrhoea in low- and middle-income countries (LMIC), are increasingly antimicrobial resistant and have no licensed vaccine. We performed genomic analyses of 1246 systematically collected shigellae from seven LMIC to inform control and identify factors that could limit the effectiveness of current approaches. We found that *S. sonnei* contributes ≥20-fold more disease than other *Shigella* species relative to its genomic diversity and highlight existing diversity and adaptative capacity among *S. flexneri* that may generate vaccine escape variants in <6 months. Furthermore, we show convergent evolution of resistance against the current recommended antimicrobial among shigellae. This demonstrates the urgent need to integrate existing genomic diversity into vaccine and treatment plans for *Shigella*, and other pathogens.

## Introduction

Shigellosis is a diarrhoeal disease responsible for approximately 212,000 annual deaths and accounting for 13.2% of all diarrhoeal deaths globally (1). The Global Enteric Multicenter Study (GEMS) was a large case-control study conducted between 2007 and 2011, investigating the aetiology and burden of moderate-to-severe diarrhoea (MSD) in children less than five years old in low- and middle-income countries (LMICs) (2). GEMS revealed shigellosis as the leading bacterial cause of diarrhoeal illness in children, who represent a major target group for vaccination (3). The aetiological agents are *Shigella*, a Gram-negative genus comprised of *S. flexneri*, *S. sonnei*, *S. boydii* and *S. dysenteriae*, with the former two serotypes causing the majority (90%) of attributable shigellosis in children in LMICs (3). Currently, the disease is primarily managed through supportive care and antimicrobial therapy. However, there has been an increase in antimicrobial resistance (AMR) among *Shigella* (4). Particularly concerning is the rise of resistance against the fluoroquinolone antimicrobial ciprofloxacin, the current World Health Organisation (WHO) recommended treatment, such that fluoroquinolone-resistant (FQR) *Shigella* is one of a dozen pathogens for which WHO notes new antimicrobial therapies are urgently needed (5). The high disease burden and increasing AMR of *Shigella* call for improvements in treatment and management options for shigellosis, and significant momentum has built to rise to this challenge.

However, there is still no licenced vaccine available for *Shigella* and one of the main challenges in its development is the considerable genomic and phenotypic diversity of the organisms (6). The distinct lipopolysaccharide O-antigen structures of *Shigella* determine its serotype and is responsible for conferring the short to medium term serotype-specific immunity following infection (7–10). Hence, considerable efforts are focused on generating O-antigen specific vaccines. However, with the exception of the single serotype *S. sonnei*, each species encompasses multiple diverse serotypes: 14 serotypes/subserotypes for *S. flexneri*, 19 for *S. boydii* and 15 for *S. dysenteriae* (11). Thus, for serotype-targeted vaccine approaches, multivalent vaccines are proposed to provide broad protection against disease (6, 12). Furthermore, while O-antigen conjugates are a leading strategy, challenge studies have recently demonstrated poor efficacy (13, 14). An attractive alternative and/or complement to serotype-targeted vaccine formulations are specific subunit vaccines which target highly conserved proteins and may offer broad protection. There are several candidates in development that have demonstrated protection in animal models (15, 16), but the degree of antigenic variation for these targets among the global *Shigella* population remains unknown.

Whole-genome sequencing analysis (WGSA) provides sufficient discriminatory power to resolve phylogenetic relationships and characterise diversity of bacterial pathogens, essential to informing vaccine development and other aspects of disease control (17, 18). However, these critical analysis tools are yet to be applied to a pathogen collection appropriate for broadly informing shigellosis control in the critical demographic of children in LMICs. Here, we apply WGSA to *Shigella* isolates sampled during GEMS, representing 1,246 systematically collected isolates from across seven nations in sub-Saharan Africa and South Asia with some of the highest childhood mortality rates (2, 19). We found evidence of the potential benefit of genomic subtype-based targeting, characterised pathogen features that will complicate current vaccine approaches, and highlighted regional differences among *Shigella* diversity, as well as determinants of AMR, including convergent evolution toward resistance against currently recommended treatments. Our analysis of this unparalleled pathogen collection informs the control and prevention of shigellosis in those populations most vulnerable to disease.

## Results and Discussion

### Regional diversity of Shigella spp. across LIMC

To date, this is the largest representative dataset of *Shigella* genomes from LMICs (*n*=1246), collected across seven sites from Asia, West Africa and East Africa, comprised of 806 *S. flexneri*, 305 *S. sonnei*, 75 *S. boydii* and 60 *S. dysenteriae* (Fig. 1A). To compare the genomic diversity of *Shigella* species, we determined the distributions of pairwise single-nucleotide polymorphism (SNP) distances and scaled the total detected SNPs against the length of the chromosome (in kbp) for each species (Fig. 1B). This revealed that *S. boydii* contained the greatest diversity (24.2 SNPs/kbp), followed by *S. flexneri* (19.5 SNPs/kbp) and *S. dysenteriae* (11.8 SNPs/kbp), with *S. sonnei* being >9.8-fold less diverse (1.2 SNPs/kbp) or >13.1-fold less diverse (0.9 SNPs/kbp) excluding two outliers (see below, Fig. 1B). This revealed that *S. sonnei* caused between 20 and 25-fold more disease relative to genomic diversity than *S. flexneri* and either *S. dysenteriae* or *S. boydii* (Fig. 1B), indicating the value of vaccination against *S. sonnei* as a comparatively conserved target relative to disease burden. Examination of the gene repertoire revealed that this relative chromosomal diversity was consistent with the accessory genome variation among species (fig. S1).

**Fig. 1.**
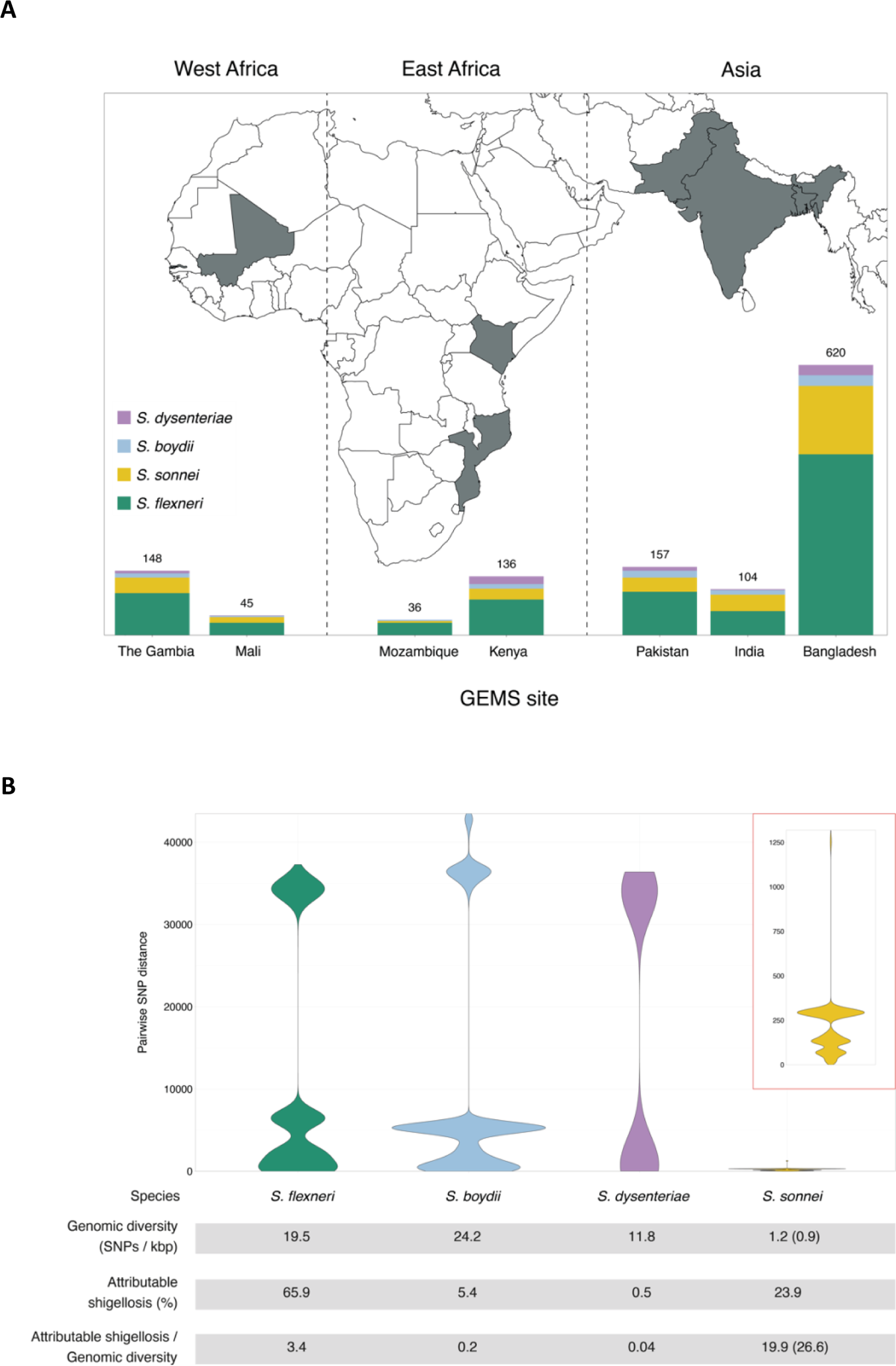
The diversity of *Shigella* spp. across seven LMIC. (**A**) Stacked bar graphs illustrate the number of isolates from each *Shigella spp.* sequenced from GEMS and used in the current study, grouped by study sites. (**B**) Pairwise genomic distances (in SNPs) among *Shigella* isolates within subgroups are shown as violin plots. A magnified plot for *S. sonnei* is displayed inside the red box. The table below the plots demonstrates for each species the genomic diversity (as measure by total number of SNPs per kbp [methods]), the contribution to GEMS shigellosis burden and the shigellosis burden relative to genomic diversity. For *S. sonnei,* the genomic diversity and shigellosis burden relative to genomic diversity that was calculated excluding the two outliers are shown in bracket.

Early global population structure studies revealed that each *Shigella* species is delineated into multiple WGSA subtypes (20–23). Specifically, *S. flexneri* is comprised of seven phylogroups (PGs) (20) and *S. sonnei* of five lineages (24). To describe the genomic epidemiology of the GEMS *Shigella* within existing frameworks we constructed species phylogenetic trees and integrated these with epidemiological metadata and publicly available genomes. The *S. flexneri* phylogeny revealed two distinct lineages separated by ∼34,000 SNPs; one comprising five previously described PGs (20) and a distant clade comprised largely of *S. flexneri* serotype 6 isolates (herein termed Sf6), contributing distinctly to the disease burden of each country (Fig. 2 and fig. S2). Phylogenetic analysis of *S. sonnei* revealed that all but two isolates belonged to the globally dominant multidrug resistant (MDR) Lineage III (21) (fig. S3). For *S. boydii* and *S. dysenteriae*, a total of four and two previously described phylogenetic clades (23, 25) were identified, respectively (fig. S4). Marked phylogenetic association of isolates with country of origin prompted an examination of species genomic diversity by region (East Africa, West Africa and Asia) and revealed that while *S. flexneri* diversity was comparable across regions, diversity varied by region for the remaining species (fig. S5). Specifically, *S. sonnei* was more genomically diverse in East Africa owing to the presence of two Lineage II isolates from Mozambique. For *S. boydii*, Asia contained greater diversity than African regions, owing to isolates belonging to additional clades. *S. dysenteriae* diversity was lower in West Africa relative to other regions by virtue of having only one circulating clade. These geographical differences highlight the importance of considering regional variations during vaccine development and that vaccine candidates should be evaluated across multiple regions.

**Fig. 2.**
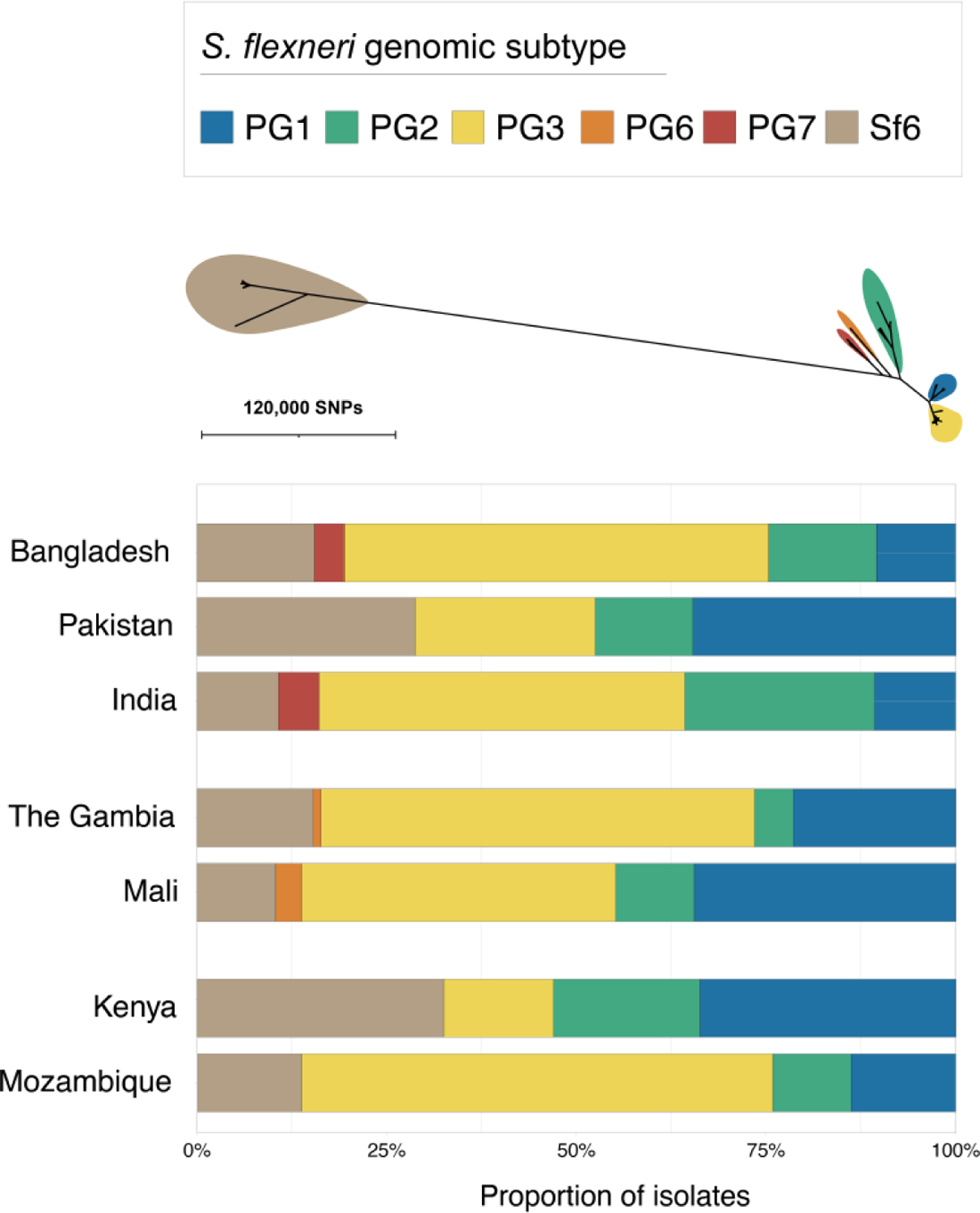
The diversity of *S. flexneri* genomic subtypes across seven GEMS study sites. An unrooted ML phylogenetic tree of *S. flexneri* genomes identified six distinct genomic subtypes, each highlighted in a different colour according to the inlaid key displayed above the tree. The bar plot below the tree demonstrates the relative frequencies of the subtypes at each study site.

### Genomic subgroups as an alternative targeting method

To explore the utility of vaccination targeting genomic subtype (relative to targeting serotype) for *S. flexneri*, we determined the relative effect size of the dominant subtype on the epidemiological outcome of shigellosis (i.e., isolates derived from case patients rather than from controls, as defined in GEMS). The dominant genomic subtype was PG3, which comprised the majority (47%, 378/806) of total isolates, as well as case (50%, 341/687) isolates, with some regional variation (Fig. 2). This resulted in an increased odds of cases (OR = 2.3, 95% CI = 1.5-3.6, *p* = 0.0001) for PG3 compared with other genomic subtypes (PGs and Sf6) (methods, table S3). The association of cases with the dominant serotype, *S. flexneri* serotype 2a (accounting for 29% (234/806) of total isolates and 31% (210/687) of case isolates) also resulted in an increased odds of cases (OR = 1.9, 95% CI = 1.7-3.2, *p* = 0.0099) (table S3). But the higher prevalence and larger effect size of PG3 relative to serotype 2a on case status offers compelling evidence that targeting vaccination by phylogroup might offer broader coverage per licenced vaccine relative to, or in combination with, a serotype-specific approach.

### Diversity of S. flexneri relevant to serotype-targeted vaccines

The development of serotype-targeted vaccines is complicated by the diversity and distribution of serotypes, which are heterogenous over time and place (8, 19, 26, 27). Furthermore, genetic determinants of O-antigen modification are often encoded on mobile genetic elements (28, 29) that can move horizontally among bacterial populations, causing the recognised, but poorly quantified phenomenon of serotype switching (20, 27, 28), which may result in the rapid escape of infection induced immunity against homologous serotypes. For our analyses of serotype switching, we focused on *S. flexneri* owing to high disease burden and serotypic diversity. Phenotypic serotyping data were overlaid onto the phylogeny and revealed that while generally there was a strong association of genotype (i.e. PG/Sf6) with serotype (Fisher’s exact test; *p*<2.20E-16), multiple serotypes were observed for each genotype (Fig. 3). The greatest serotype diversity was observed in PG3, comprised of seven distinct serotypes and two subserotypes. Correlation of serotypic diversity (number of serotypes) and genomic diversity (maximum pairwise SNP distance within genotype) revealed no evidence for an association, but a significant positive correlation of serotypic diversity with the number of isolates in each genotype was found (fig. S6), indicating that serotype diversity scales with prevalence.

**Fig. 3.**
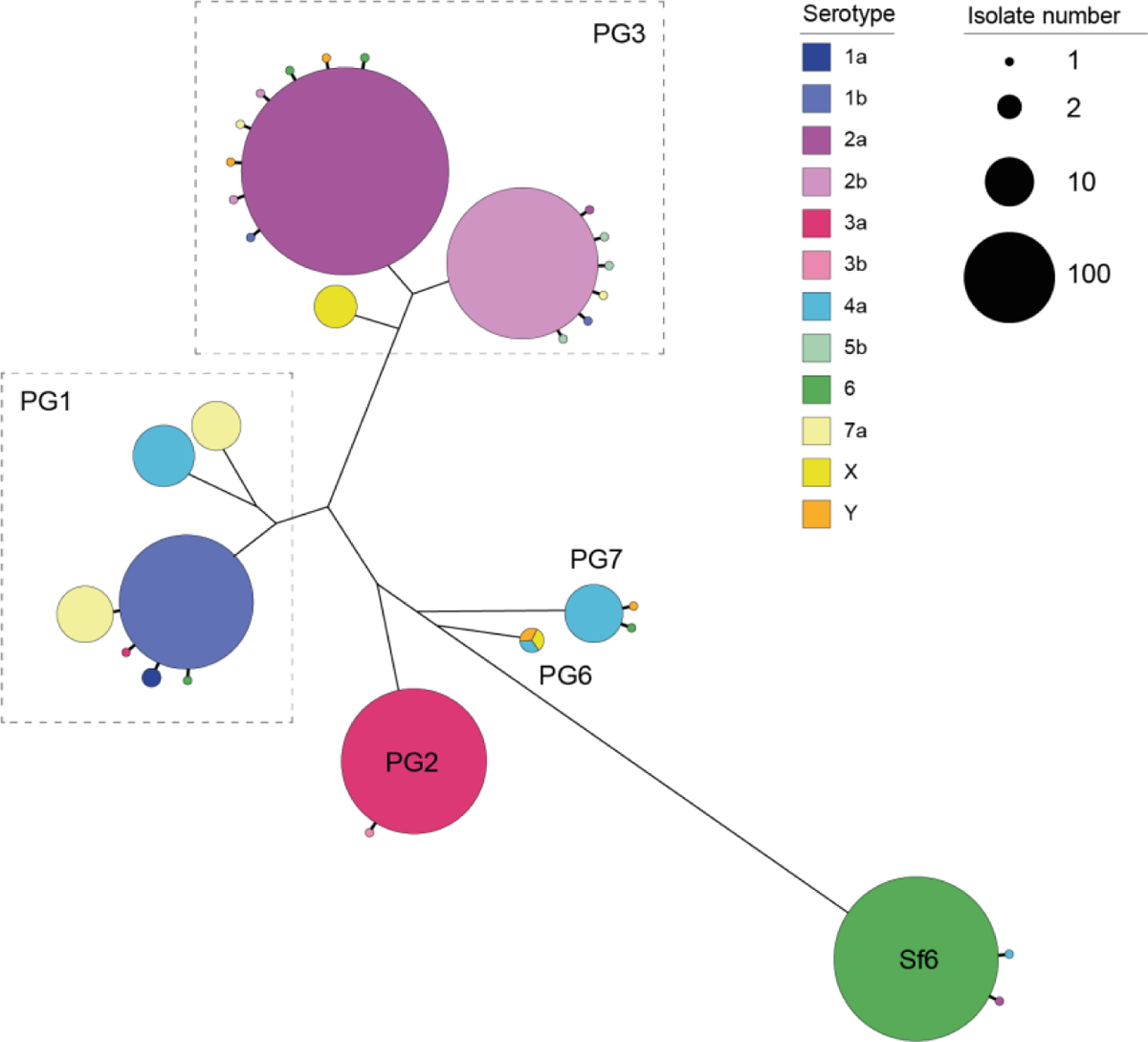
Diversity of *S. flexneri* population with respect to serotype switching. The unrooted *S. flexneri* phylogenetic tree is shown with the five phylogroups (PG1-PG7) and Sf6 labeled accordingly. For each genomic subtype, monophyletic clusters of the dominant serotype are shown collapsed into bubbles coloured according to the inlaid key. Single isolates or groups of isolates within a subtype of an alternative serotype are represented by further branches, indicating a single serotype switch. The number of isolates within a single cluster is represented through bubble size.

To qualitatively and quantitatively determine serotype switching across *S. flexneri,* we examined the number of switches occurring within each genotype. A switching event was inferred when a serotype emerged (either as a singleton or monophyletic clade) that was distinct from the majority (>65%) serotype within a genotype (Fig. 3 and fig. S7). PG6 was excluded from the analysis, as only three isolates from GEMS belonged to this genotype and a dominant serotype could not be inferred. Quantitatively, this revealed serotype switching was infrequent, with only 26 independent switches (3.3% of isolates) identified across the five *S. flexneri* genotypes. Although the frequency of switching varied across the genotypes, statistical support for an association of serotype switching with genotype fell short of significance (Fisher’s exact test; *p* = 0.09). Qualitatively, the majority (22/26) of switching resulted in a change of serotype, with few (4/26) resulting in a change of subserotype. Examination of O-antigen modification genes revealed that serotype switching was facilitated by changes in the composition of phage-encoded *gtr* and *oac* genes in the genomes, as well as point mutations in these genes (table S4). Our data also revealed that few (4/26) switching events resulted in more than two descendant isolates (fig. S7). This indicates that while natural immunity drives the fixation of relatively few serotype-switched variants in the short term, the potential pool of variants that could be driven to fixation by vaccine-induced selective pressure following a serotype-targeted vaccination program is much larger.

In order to estimate the likely timeframe over which serotype switching events might be expected to occur, we estimated the divergence time of the phylogenetic branch giving rise to each switching event. To streamline the analysis, we focused on two subclades of PG3, the most prevalent phylogroup, in which seven independent serotype switching events were detected (fig. S8). Based on the timeframes observed within our sample (spanning 4 years from 2007 to 2010), serotype switching was estimated to occur within an average of 348 days, ranging from 159 days (95% highest posterior density [HPD]: 16 - 344) to 10206 days (28 years) (95% HPD: 5494 - 15408) (table S5). Taken together, our data shows that although serotype-switching frequency is low, it can occur over relatively short timeframes and lead to serotype replacement such that non-vaccine serotypes could replace vaccine serotypes following a vaccination program, as has been observed for *Streptococcus pneumoniae* (30, 31). These elucidated serotype switching dynamics (i.e. switching occurring over short timeframes and quantitatively proportional to disease burden) highlights the value of a multivalent vaccine and geographically coordinated implementation of *Shigella* vaccination.

### Heterogeneity among Shigella vaccine protein antigens

Conserved antigen-targeted vaccines can overcome some hurdles of serotype-targeted vaccines. Hence, we performed detailed examination of six protein antigens that are currently in development and have demonstrated protection in animal models (Table 1). First, we assessed the distribution of the candidates among GEMS *Shigella* isolates which revealed that the proportional presence of antigens varied across species and with genetic context. Specifically, genes encoded on the virulence plasmid (*ipaB*, *ipaC*, *ipaD*, *icsP*) were present in >85% of genomes for each species with the exception of *S. sonnei* (fig. S9). The low proportion (≤5%) of virulence plasmid encoded genes detected among *S. sonnei* was caused by a similarly low detection of the virulence plasmid among *S. sonnei* (6%), which likely arose due to loss during sub-culture (32). In contrast, the chromosomally encoded *ompA* was present in >98% of all isolates, while the *sigA* gene (carried on the chromosomally integrated SHI-1 pathogenicity island (17)) was present in 99% of *S. sonnei* genomes, but only 63% of *S. flexneri* genomes. Notably, among *S. flexneri* genomes, the *sigA* gene was exclusively found in PG3 and Sf6, and present in >96% of isolates in each genotype) (fig. S2), indicating an appropriate distribution for targeting the two genotypes. Second, we assessed the antigens for amino acid variation and modelled the likely impact of detected variants, as antigen variation may also lead to vaccine escape, as demonstrated for the P1 variant of SARS-CoV2 (33, 34). We determined the distribution of pairwise amino acid (aa) sequence identities per antigen against *S. flexneri* vaccine strains for each species (methods). Overall, sequence identities were >90% but varied with antigen (fig. S9). For example, OmpA was present in the highest proportion of genomes, but showed ∼5% sequence divergence, while SigA was present in fewer genomes, but exhibited little divergence (<0.5%) among species. The least conserved sequence was IpaD, ranging from 3 to 7% divergence within species.

**Table 1.** *Shigella* antigen vaccine candidates examined in the current study.

| Vaccine candidate | Development stage | Location | Reference |
| --- | --- | --- | --- |
| IcsP (OmpP) | Preclinical | Virulence plasmid | Czerkinsky and Kim (37) |
| SigA | Preclinical | Chromosome (pathogenicity island) | Czerkinsky and Kim (37) |
| IpaB | Phase I | Virulence plasmid | Martinez-Beccera (16); Riddle et al (83); Tribble et al (84) |
| IpaC |  |  |  |
| IpaD |  |  |  |
| OmpA | Preclinical | Chromosome | Pore *et al* (85) |

Not all antigenic variation will affect antibody binding, so we performed *in silico* analyses of the detected variants to assess whether they may compromise the antigens as vaccine targets. Again, we focused our analyses on *S. flexneri* owing to its high disease burden and the likely complication of serotype-based vaccination strategies for this species. We detected 121 variants across the six antigens, the majority (79%) of which correlated with genotype (i.e. belonging to either PGs 1-5 or Sf6, fig. S11). We then determined if amino acid variants were located in immunogenic regions (i.e. epitope/peptide fragment) (fig. S10) and assessed their potential destabilization of protein structure through *in silico* protein modelling. For IpaB, IpaC and IpaD, the epitopes have been empirically determined (35, 36). The sequence and location of peptide fragments of SigA, IcsP and OmpA used in vaccine development are available (37, 38). Variants located within the immunogenic regions were identified for all antigens relative to PG3 reference sequences (methods, Fig. 4). Only 4 of 121 variants were predicted to be highly destabilising to protein structure, and these occurred in: OmpA (residue 89) at a periplasmic turn, SigA (residues 1233 and 1271) in adjacent extracellular turns in the translocator domain (fig. S12), and in IpaD (residue 247) within a beta-turn-beta motif flanking the intramolecular coiled-coil (Fig. 4). While it remains possible that these mutations could affect antigenicity through the disruption of folding or global stability, it is less likely than if they occurred in immunogenic regions. These results thus indicate that it is less likely that existing natural variation will compromise antigen-based vaccine candidates for *Shigella* compared with serotype-based vaccines. However, our approach is limited and the knowledge base incomplete. For example, there was no suitable template available for IpaC, and some epitopes were predicted to be in membrane regions which should be inaccessible to antibodies, indicating the need for more accurate publicly available protein structures to be developed for many of the vaccine antigen candidates.

**Fig. 4.**
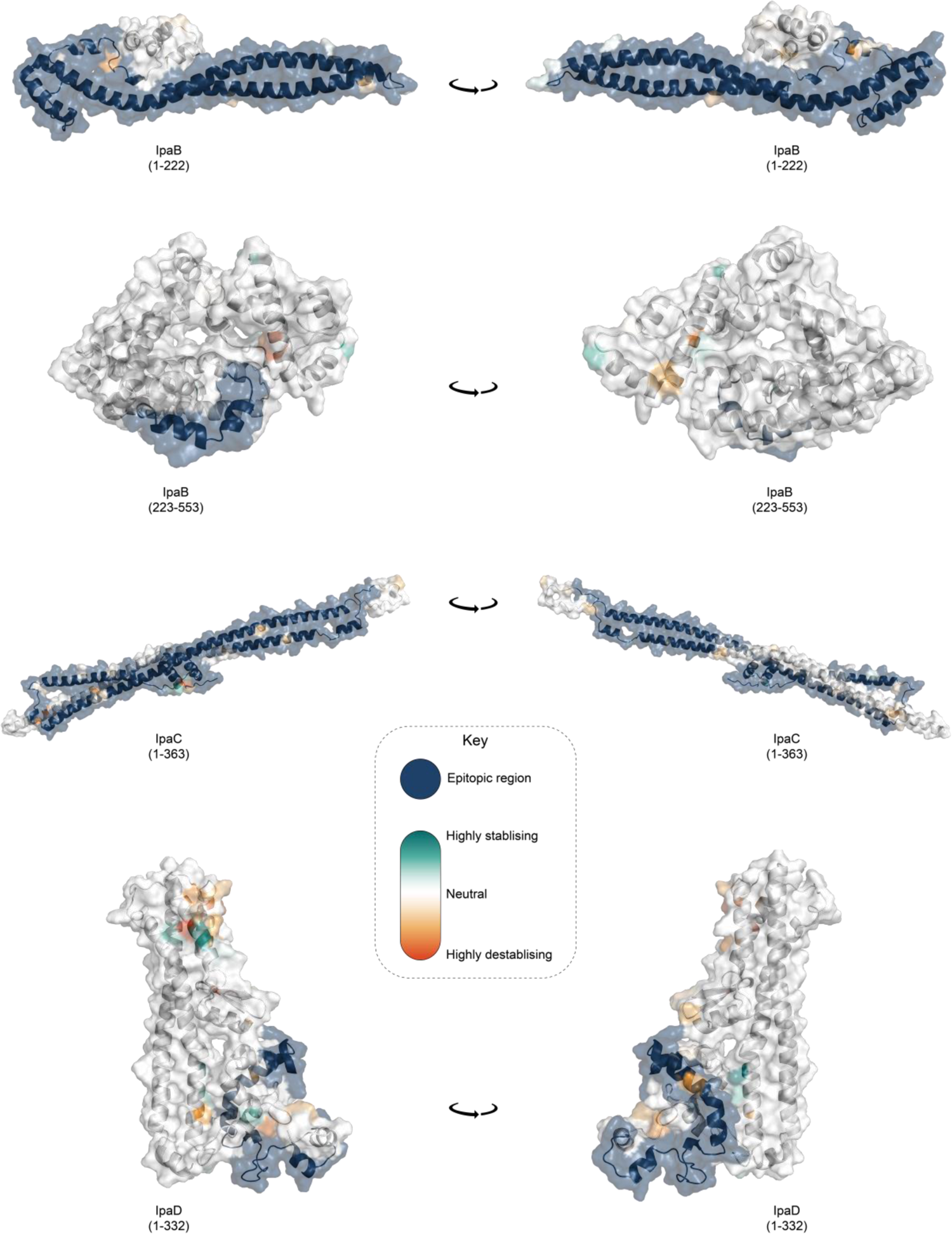
Visualization of mutations and its predicted effect on modeled IpaB, IpaC and IpaD protein antigens. Visualisation of mutations on modelled proteins IpaB, IpaC and IpaD. The protein residue ranges modelled are shown in brackets. Blue region represents empirically determined epitopes. Mutations identified within the proteins are coloured using the scale shown in the inlaid key, where highly destabilising mutations are dark orange and highly stabilising mutations are dark green.

### Region-specific details of antimicrobials as a stop gap

**Until a licensed vaccine is available, we must continue to treat shigellosis with supportive care and antimicrobials, for which the current WHO recommendation is the fluoroquinolone, ciprofloxacin** (**39**). However, FQR *Shigella* is currently on the rise and spreading globally (40). To examine AMR prevalence among GEMS isolates for evaluating treatment recommendations, we screened for known genetic determinants (horizontally acquired genes and point mutations) conferring resistance or reduced susceptibility to antimicrobials. Although we used only minimal phenotypic data, phenotypic resistance and genotypic prediction correlate well in *S. flexneri* and *S. sonnei* (41, 42). Our analysis revealed that 95% (1189/1246) of isolates were multidrug resistant (MDR), carrying AMR determinants against three or more antimicrobial classes (Fig 5A). *S. flexneri* exhibited the greatest diversity of AMR determinants, with a total of 45 identified determinants across the population, comprising of 38 AMR genes and 7 point mutations (fig. S13 and table S1), and an extensive AMR genotype diversity of 72 unique resistance profiles (Fig. 5A and fig. S14). In contrast, *S. sonnei* exhibited the least diversity, with only 23 AMR determinants and 21 unique resistance profiles. An intermediate and comparable degree of AMR diversity was observed for both *S. dysenteriae* and *S. boydii*.

**Fig. 5.**
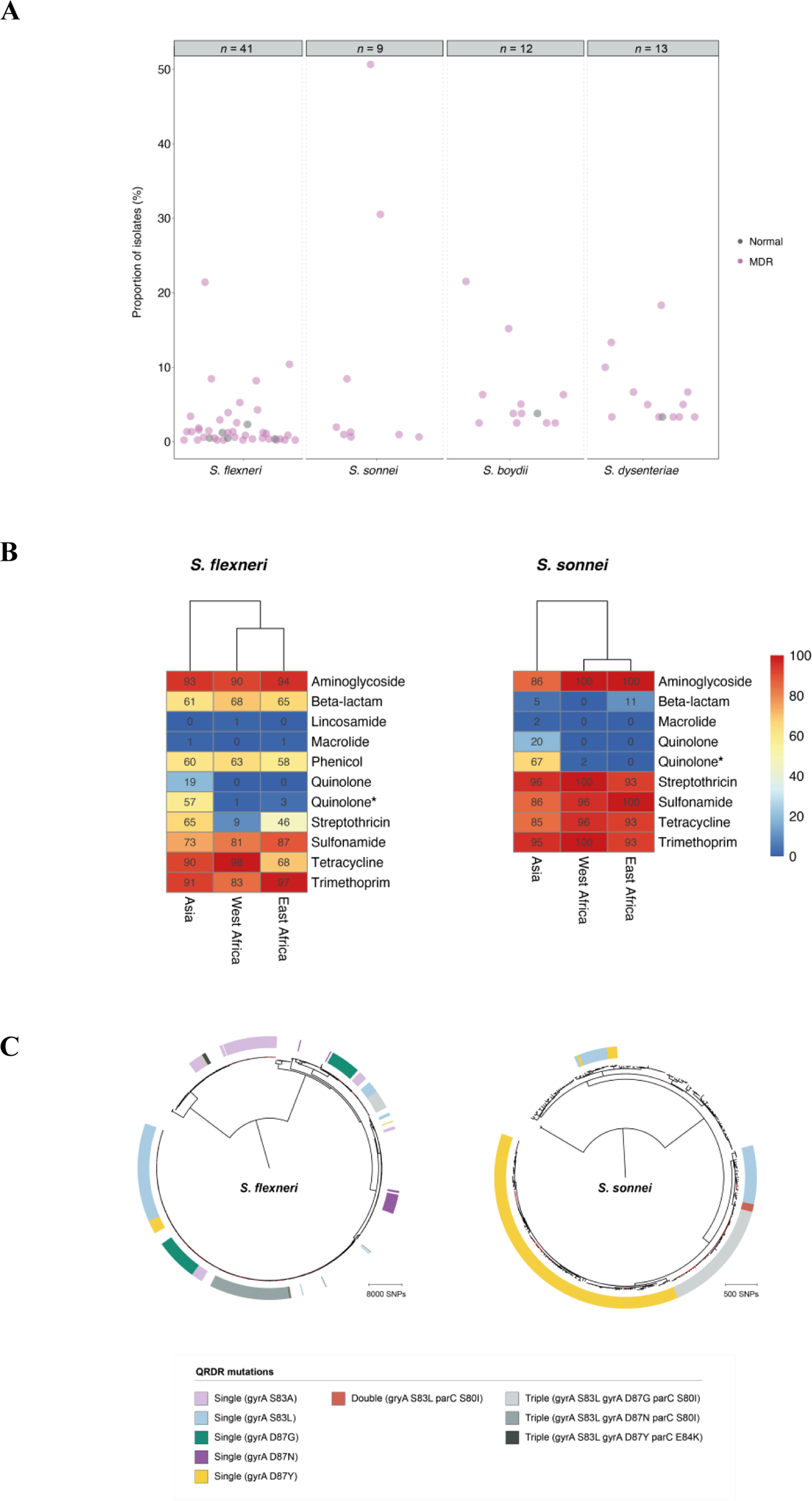
AMR genotypic profile diversity and convergent evolution of ciprofloxacin resistance. (**A**) Frequencies of AMR genotypic profiles among *Shigella* spp. Each point in the scatterplot represents a unique AMR genotype profile: the proportion of isolates with a particular profile is displayed along the y-axis. Profiles identified in only a single isolate are not displayed. MDR genotypic profile conferring resistance or reduced suppressibility to three or more drug classes are highlighted in pink, and normal AMR genotype profile conferring resistance or reduced suppressibility in fewer than three drug classes are in grey. Numbers displayed above the plot represents the number of AMR genotype profiles plotted for each species. (**B**) Detection of known AMR genetic determinants associated with drug class grouped by country. Each cell in the heatmap represents the percentage of isolates from a region containing genetic determinants associated with resistance to a drug class. Genetic determinant conferring reduced susceptibility to quinolone is indicated with an asterisk. (**C**) The genetic convergent evolution of ciprofloxacin resistance in *S. flexneri* and *S. sonnei*. The presence of multiple monophyletic clades of QRDR mutations (single, double, or triple according to the inlaid key) conferring reduced susceptibility or resistance to ciprofloxacin is shown in the outer ring. B and C for *S. boydii* and *S. dysenteriae* are shown elsewhere (fig. S15).

Overall, a high frequency of AMR genes conferring resistance against aminoglycoside, tetracycline, trimethoprim, and sulphonamide antimicrobials was observed, while resistance against other antimicrobial classes varied with region and species (Fig. 5B). The extended spectrum beta-lactamase gene *blaCTX-M-15* was detected in a small (9/1246) percentage of isolates, and genes conferring resistance to macrolides and lincosamides were also infrequent (fig. S13), indicating that the recommended second-line treatments likely remain effective antimicrobials (43).

However, higher rates of resistance were found against the first-line treatment. FQR in *Shigella* can be conferred through the acquisition of FQR-genes or, more typically, by point mutations in the chromosomal Quinolone Resistance Determining Region (QRDR) within the DNA gyrase (*gryA*) and the topoisomerase IV (*parC*) genes. Single and double QRDR mutations are known to confer reduced susceptibility to ciprofloxacin and are evolutionary intermediates on the path to resistance, conferred by triple mutations in this region (41, 44). Overall, FQR-genes were uncommon in *S. flexneri* (4%, 33/806), *S. sonnei* (1%, 3/305) and *S. dysenteriae* (7%, 4/60), but were present in 32% (24/75) of *S. boydii*. QRDR mutations were identified in all species (fig. S13), but were more common among *S. sonnei* (65%, 199/305) and *S. flexneri* (54%, 435/806) than compared with *S. boydii* (15%, 11/75) and *S. dysenteriae* (30%, 18/60). Among these, triple QRDR mutations were identified in 13% (106/806) of *S. flexneri* and 14% (44/305) of *S. sonnei*. Analysis of the QRDR mutants across the phylogenies indicate marked convergent evolution toward resistance across the genus. Specifically, all triple QRDR mutant *S. sonnei* belonged to one monophyletic subtype (previously described as globally emerging from Southeast Asia (45)), while three distinct triple QRDR mutational profiles were found across three polyphyletic *S. flexneri* genotypes (Fig. 5C). Thus, the polyphyletic distribution of single, double, and triple QRDR mutants indicates continued convergent evolution of lineages with reduced susceptibility or resistant to FQR.

We then stratified the dataset by geographic region which revealed that FQR were largely associated with isolates from Asia where fluoroquinolones are more frequently used compared to African sites (Fig. 5B) (46), which is consistent with trends observed in atypical enteropathogenic *Escherichia coli* isolated from GEMS (46). Our analyses thus suggest that for the period of GEMS trial (2007 – 2011), 17% (150/881) of *Shigella* isolates from Asia were resistant and 58% (508/881) had reduced susceptibility to the WHO recommended antimicrobial. The high level of reduced susceptibility together with marked convergent evolution toward resistance suggests that management of shigellosis with fluroquinolones at these sites may soon be ineffective and regional antimicrobial treatment guidelines may require updating. These results indicate the value of AMR and microbiological surveillance in LMICs and the control and management of shigellosis will be improved by initiatives such as the Africa Pathogen Genomics Initiative (47) and the WHO Global Antimicrobial Resistance Surveillance System (48).

## Conclusions

Pathogen genomics is a powerful tool that has a wide range of applications to help combat infectious diseases. Here, we have applied this tool to an unparalleled systematically collected *Shigella* dataset to characterise the relevant population diversity of this pathogen across LMICs in a pre-vaccine era. Our results revealed that current antimicrobial treatment guidelines for shigellosis should be updated, and that improved surveillance will be essential to guide **antimicrobial stewardship**. This study has also highlighted the urgent need to continue the development of *Shigella* vaccines for children in endemic areas. The genomic diversity in *Shigella* presents a major hurdle in controlling the disease and we have demonstrated the anticipated pitfalls of current vaccination approaches, emphasising the importance of considering the local and global diversity of the pathogens in vaccine design and implementation. Although our results are focused on shigellosis, our approach is translatable to other bacterial pathogens which is particularly relevant as we enter the era of vaccines for AMR.

## Materials and Methods

### Dataset, bacterial isolates and sequencing

A total of 1,264 *Shigella* isolates from GEMS were under investigation in this study (2, 3). All isolates were derived from stool samples/rectal swabs: their identification, confirmation and isolation have been described previously (19). A total of 1,344 isolates were sequenced at the Earlham institute, with genomic DNA extraction, sequencing library construction and whole genome sequencing carried out according to the Low Input Transposase Enabled (LITE) pipeline described by Perez-Sepulveda *et al* (49). Among these, 225 isolates failed QC with a mean sample depth of coverage <10x and an assembly size of <4MB and were re-sequenced. For these isolates, genomic DNA was re-extracted at the University of Maryland School of Medicine (Baltimore, Maryland) from cultures grown in Lysogeny Broth overnight. DNA was extracted in 96-well format from 100 μL of sample using the MagAttract PowerMicrobiome DNA/RNA Kit (Qiagen, Hilden, Germany) automated on a Hamilton Microlab STAR robotic platform. Bead disruption was conducted on a TissueLyser II (20 Hz for 20 min) instrument in a 96 deep well plate in the presence of 200 μL phenol/chloroform. Genomic DNA was eluted in 90 μl water after magnetic bead clean up and the resulting genomic DNA was quantified by Pico Green. The genomic DNA was shipped to the Centre for Genomic Research (University of Liverpool) for whole genome sequencing. Sequencing library was constructed using NEBNext® Ultra™ II FS DNA Library Prep Kit for Illumina and sequenced on the Illumina® NovaSeq 6000 platform, generating 150bp paired-end reads. An additional 125 publicly available *Shigella* and *E. coli* reference genomes were included in the analyses. Details of GEMS and reference genomes analysed in this study are listed in table S1 and table S2, respectively.

### Sequence mapping and variant calling

Adaptors and low-quality bases were trimmed with Trimmomatic v0.38 (50), reads qualities were assessed using FastQC v0.11.6 (https://www.bioinformatics.babraham.ac.uk/projects/fastqc/) and MultiQC v1.7 (51). Filtered reads were mapped against *Shigella* reference genomes with BWA mem v0.7.17 (52) using default parameters. *S. flexneri*, *S. sonnei*, *S. boydii* and *S. dysenteriae* sequencing reads were mapped against reference genomes from Sf2a strain 301 (accession NC_004337), Ss046 (accession NC_007384), Sb strain CDC 3083-94 (accession NC_010658) and Sd197 (accession NC_007606), respectively. Mappings were filtered and sorted using the SAMtools suite v1.9-47 (53), and optical duplicate reads were marked using Picard v2.21.1-SNAPSHOT MarkDuplicates (http://broadinstitute.github.io/picard/). QualiMap v2.2.2 (54) was used to evaluate mapping qualities and estimate mean sample depth of coverage. Sequencing reads for isolates sequenced using the LITE pipeline and re-sequenced at CGR were combined to increase overall sample depth of coverage. Sequence variants were identified against reference using SAMtools v1.9-47 mpileup and bcftools v1.9-80 (53). Low quality SNPs were filtered if mapping quality <60, Phred-scaled quality score <30 and read depth <4.

### Phylogenetic reconstruction and inference of genomic diversity

Filtered SNP variants were used to generate a reference-based pseudogenome for each sample, where regions with depth of coverage >4x were masked in the pseudogenome. Additionally, regions containing phage (identified using PHASTER (55)) and insertion sequences were identified from the reference genomes, and co-ordinates were used to mask these sites on the pseudogenomes using BEDTools v2.28.0 maskfasta (56). For each species, chromosome sequences from the masked pseudogenomes were extracted and concatenated. Gubbins v2.3.4 (57) was used to remove regions of recombination and invariant sites from the concatenated pseudogenomes. This generated a chromosomal SNP alignment length of 78,251 bp for *S. flexneri* (*n*=806), 5,081 bp for *S. sonnei* (*n*=305), 98,842 bp for *S. boydii* (*n*=75) and 45,031 bp for *S. dysenteriae* (*n*=60). Maximum-likelihood phylogenetic reconstruction was performed independently for each species and inferred with IQ-TREE v2.0-rc2 (58) using the FreeRate nucleotide substitution, invariable site and ascertainment bias correction model, with 1000 bootstrap replicates. In order to contextualise GEMS isolates within the established genomic subtypes and to infer the most appropriate root for each species tree, phylogenetic trees were reconstructed including publicly available reference genomes of isolates from previously defined lineages/phylogroups/clades and *E. coli* isolates (table S2). Phylogenetic tree for *S. flexneri*, *S. boydii* and *S. dysenteriae* was rooted using *E. coli* strain IAI1-117 (accession SRR2169557) as an outgroup, respectively. Phylogenetic tree for *S. sonnei* was midpoint rooted. Visualizations were performed using interactive Tree of Life (iTOL) v6.1.1 (59).

To measure the extent of *shigella* genomic diversity among GEMS population, pairwise SNP distance was determined from the alignment of core genome SNPs identified outside regions of recombination using snp-dists v0.7.0 (https://github.com/tseemann/snp-dists). For each species, the genomic diversity, measured by SNPs per kbp, was determined by dividing the core genome SNP alignment length by the core genome size (*S. flexneri* 4,015,307 bp, *S. sonnei* 4,177,070 bp, *S. boydii* 4,088,693 bp and *S. dysenteriae* 3,821,602 bp). Scaling the proportion of disease burden attributable by the genome diversity of each species, the percentage of species contribution to GEMS shigellosis disease burden was divided by the number of SNPs per kbp.

### Serotype switching time frame inference

To estimate the likely time frame of serotype switching, we performed temporal phylogenetic reconstruction in order to infer the time of divergence along branches exhibiting serotype switching. We streamlined the analysis and focused on isolates belonging to two subclades of *S. flexneri* PG3. First, for each of the two subclades (*n*=99 and *n*=45), a maximum-likelihood phylogeny was reconstructed based on genome multiple sequence alignments (described above). Then, TempEst v1.5.3 (60) was used determine if there is sufficient temporal signal in the data by inferring linear relationship between root-to-tip distances of the phylogenetic branches with the year of sample isolation. Data from both subclades revealed positive correlation between sampling time and phylogenetic root-to-tip divergence, with R^2^ of 0.186 and 0.111 (fig. S16). Once temporal signals within each of the two datasets were confirmed, core genome SNP alignments of length 559 bp and 1,244 bp were analysed independently using BEAST2 v2.6.1 (61). The parameters were as follows: dates specified as days, bModelTest (62) implemented in BEAST2 was used to infer the most appropriate substitution model, a relaxed log normal clock rate with a coalescent Bayesian skyline model for population growth. A total of five independent chains were performed, each with chain length of 250,000,000, logging every 1,000 and accounting for invariant sites. Convergence of each run was visually assessed with Tracer v1.7.1 (63), with all parameter effective sampling sizes ≥200. Tree files were sampled and combined using LogCombiner v2.6.1, the combined files were then summarised using TreeAnnotator v2.6.0 with 10% burn-in to generate Maximum Clade Credibility tree (64). Divergence time was inferred by reading the branch length from the most recent common ancestor to the first sampled isolate that serotype-switched.

### Genome assembly and annotation

Draft genome sequences were assembled using Unicycler v0.4.7 (65) with –min_fasta_length set to 200. QUAST v5.0.2 (66) was used to assess the qualities of the assemblies. Assemblies with total assembly length outside the range of <4MBP and >6.4Mbp were removed. Resulting in an average length of 4,275,508 bp (range: 4 4,004,109 – 4,538,734 bp) for *S. flexneri*, 4,264,097 bp (range: 4,008,630 – 4,779,279 bp) for *S. sonnei*, 4,227,671 bp (range: 4,000,714 – 4,689,815 bp) for *S. boydii* and 4,297,921 bp (range: 4,040,642 – 4,659,860 bp) for *S. dysenteriae*. An average N50 value of 29,804 bp (range: 6,810 – 34,658 bp) was generated for *S. flexneri*, 23,961 bp (range: 11,547 – 30,008 bp) for *S. sonnei*, 20,835 bp (range: 15,323 – 40,119 bp) for *S. boydii* and 22,137 bp (range: 14,090 – 31,358 bp) for *S. dysenteriae*. Draft genomes were annotated using Prokka v1.13.3 (67).

### Pangenome analysis

The pangenome of each species was defined using Roary v3.12.0 (68) without splitting paralogues. The pangenome accumulation curves were generated separately for each species using the specaccum function from Vegan v2.5-7 (https://github.com/vegandevs/vegan/), with 100 permutations and random subsampling. Inspections of the variable gene content showed that all four species had open pangenomes, implying that the number of unique gene count increases with the addition of newly sequenced genomes.

### Shigella flexneri molecular serotyping

*Shigella* serotype data was provided by collaborators at the University of Maryland School of Medicine (Baltimore, Maryland), serotyping was performed as previously describe (19). *In silico* serotyping of *S. flexneri* genomes was performed using ShigaTyper v1.0.6 (69) which detects the presence of serotype-determining genetic elements from sequencing reads to predict serotype. ShigaTyper predictions were 84% concordant to the serotype data provided. SRST2 v2 (70) was used to detect mutations within serotype-determining genetic elements, run against ShigaTyper sequence database with default parameters.

### Protein antigen screening

To determine the presence of antigen vaccine candidates among GEMS *Shigella* isolates, genes of the antigen vaccine candidates was screened against draft genome assemblies using screen_assembly (17) with a threshold of ≥80% identity and ≥70% coverage to the reference sequence. Reference sequences for *ipaB*, *ipaC*, *ipaD* and *icsP* were derived from *S. flexneri* 5a strain M90T (accession GCA_004799585) and *ompA* and *sigA* was derived from *S. flexneri* 2a strain 2457T (accession NC_004741), both strains are commonly used in the laboratory for vaccine development. Antigen sequence variations were determined by examining the BLASTp (71) percentage identity against relevant query reference sequence. Allelic variations of antigen vaccine candidates among *S. flexneri* population were identified manually by visualising amino acid sequence alignments using AliView v1.26 (72).

### Protein antigen modelling

In order to assess the effect of point mutations on protein stability and vaccine escape, six antigen candidates from *S. flexneri* PG3 were modelled: OmpA, SigA, IcsP, IpaB, IpaC and IpaD (Table 1). PG3 was selected as it is the most prevalent phylogroup and is therefore the target of current vaccine development. To model the antigen targets, we first searched for a suitable template using HHPred (73, 74). Five of the six proteins (OmpA, SigA, IcsP, IpaB and IpaD) had suitable homologues available. To improve the performance of the comparative modelling, the signal peptides for OmpA, SigA and IcsP were removed and OmpA, SigA and IpaB were modelled in two parts to make use of optimal templates. RosettaCM (75) was used to generate 200 models for each of the five proteins using the single best available template. For IpaC, where no suitable templates were available, trRosetta (76) was used to create five de novo predicted models. The best model for each antigen candidate was selected using QMEAN’s average local score. QMEANbrane (77, 78) was used for suitable membrane proteins (IpaB, IpaC & IpaD), otherwise QMEANDisCo (77) was used (table 6). Full details of the modelling and ranking are shown in table 7. The effect of point mutations on the stability of the antigen candidates was assessed using PremPS, and the default criterion of (ΔΔG > 1 kcal mol^-1^) used to defining highly destabilising mutations (79).

### Detection of AMR genetic determinants and AMR testing

To detect the presence of known genetic determinants for AMR, AMRFinderPlus v3.9.3 (80) was used to screen draft genome assemblies against the AMRFinderPlus database, which is derived from the Pathogen Detection Reference Gene Catalog (https://www.ncbi.nlm.nih.gov/pathogens/). AMRFinderPlus was performed with the organism-specific option for *Escherichia*, to screen for both point mutations and genes, and filter out uninformative genes that were nearly universal in a group. Output was then filtered to remove genetic determinants identified with ≤80% coverage and ≤90% identity. The presence of *S. sonnei* virulence plasmid was confirmed using short-read mapping using BWA mem (as described above) against the reference virulence plasmid from Ss046 (GenBank accession CP000039.1). Presence of the plasmid was defined by mapping of >60% breadth of coverage across the reference. Visualisations of AMR resistance profiles were performed with UpSetR v2.1.3 (81). Four *S. flexneri* isolates with triple QRDR mutations were phenotypically tested for ciprofloxacin resistance using the Kirby-Bauer standardized disk diffusion method (82).

### Statistical analyses

The strength of association between *S. flexneri* genomic subtype and serotype with the occurrence of case outcome was calculated using MedCalc’s odds ratio calculator v20 (https://www.medcalc.org/calc/odds_ratio.php) to report the odds ratio, 95% confidence interval and statistical association. Association of genomic subtype with serotype and serotype switching was tested using Fisher’s exact test. Linear regression analysis was used to determine the correlation between serotype diversity to various properties of genomic subtype. Both analyses were performed using R v4.0.3.

## Acknowledgements

We acknowledge and thank members of Baker group and Lab H at the University of Liverpool, and Rodrigo Bacigalupe at KU Leuven for invaluable discussions. We also thank Jay Hinton and Blanca Perez Sepulveda for logistical support orchestrating the thermolysate shipping. The authors are grateful to Sam Haldenby, Matthew Gemmell and Richard Gregory and the Centre for Genomics Research, University of Liverpool for technical support. The authors acknowledge Dr. Irene Kasumba, Ms. Jennifer Jones, Mr. Sunil Sen and Ms. Jasnehta-Permala-Booth for preparing GEMS *Shigella* isolates for sequencing and antimicrobial testing.

## Author contributions

R.J.B performed majority of the data analysis and interpretation of the results under the scientific guidance of K.S.B. A.J.S and D.J.R performed *in silico* protein antigens modelling and prediction of the impacts of amino acid substitutions on protein stability. C.V.P supported Bayesian Evolutionary Analysis by Sampling Trees. S.M.T. prepared and provided GEMS *Shigella* isolates and metadata. DR contributed to sample preparation. R.J.B and K.S.B drafted the manuscript. All authors contributed to editing of the manuscript.

---

**Fig. S1.**
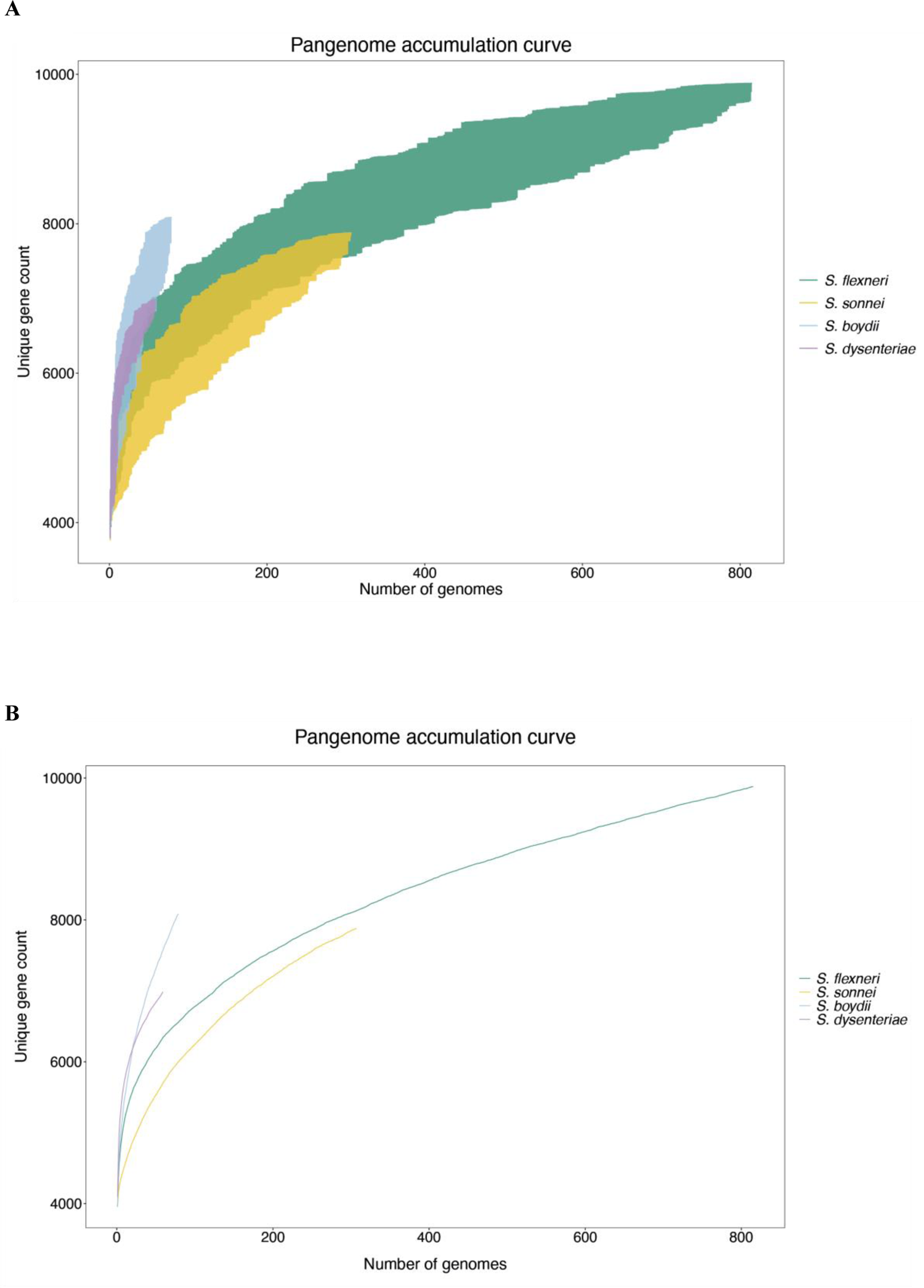
Pangenome accumulation curve of *S. flexneri*, *S. sonnei*, *S. boydii* and *S. boydi*. Each curve demonstrates the number of unique protein coding genes in the pangenome as a new genome is randomly added, with the number of genomes plotted along the x-axis. Random permutation of the data were subsampled 100 times, in which genomes are subsampled without replacement at each iteration. The y-axis shows the minimum and maximum range of unique gene count after each iteration in (A) and the mean value in (B).

**Fig. S2.**
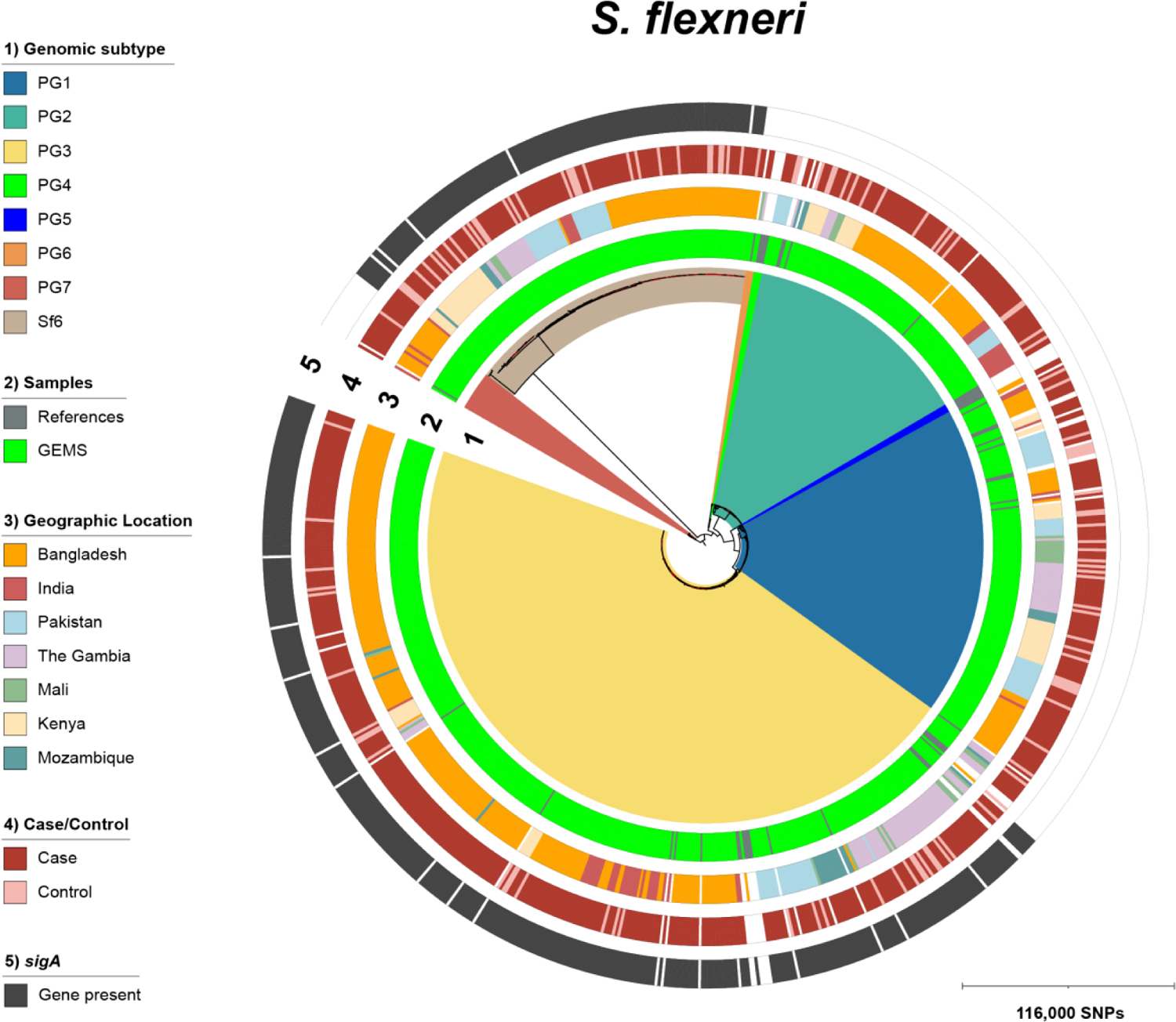
Phylogeny of *S. flexneri* population from GEMS. ML phylogenetic tree constructed using core genome SNPs from alignments of 817 *S. flexneri* genomes from GEMS and publicly available genomes. Tree was rooted using *E. coli* genome. The outer concentric rings illustrate different genotypic and epidemiological data according to the numbered inlaid keys displayed next to the tree. Scale bars represents the number of SNPs.

**Fig. S3.**
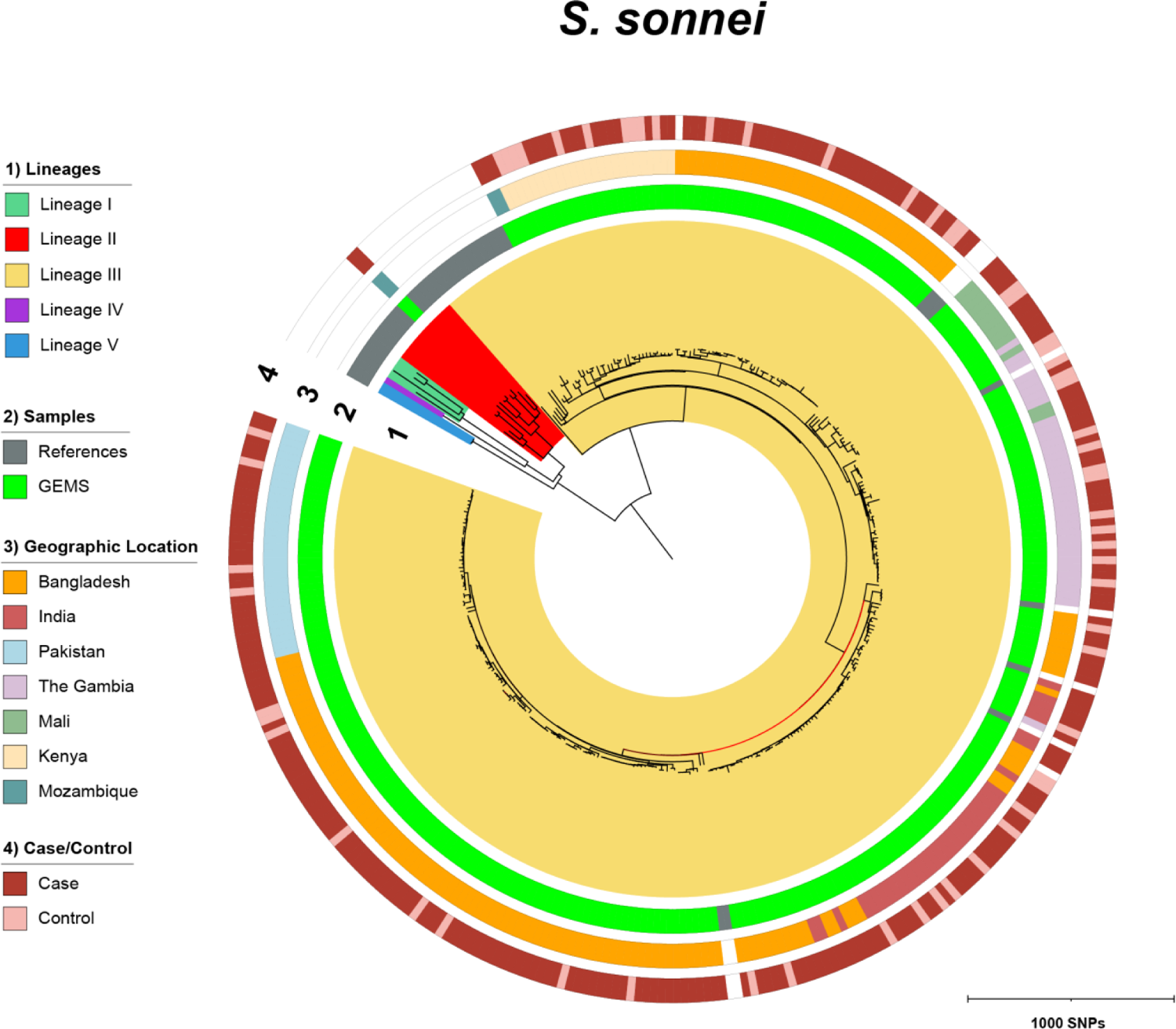
Phylogeny of *S. sonnei* population from GEMS. Midpoint rooted ML phylogenetic tree constructed using core genome SNPs from alignments of 308 *S. sonnei* genomes from GEMS and publicly available genomes.

**Fig. S4.**
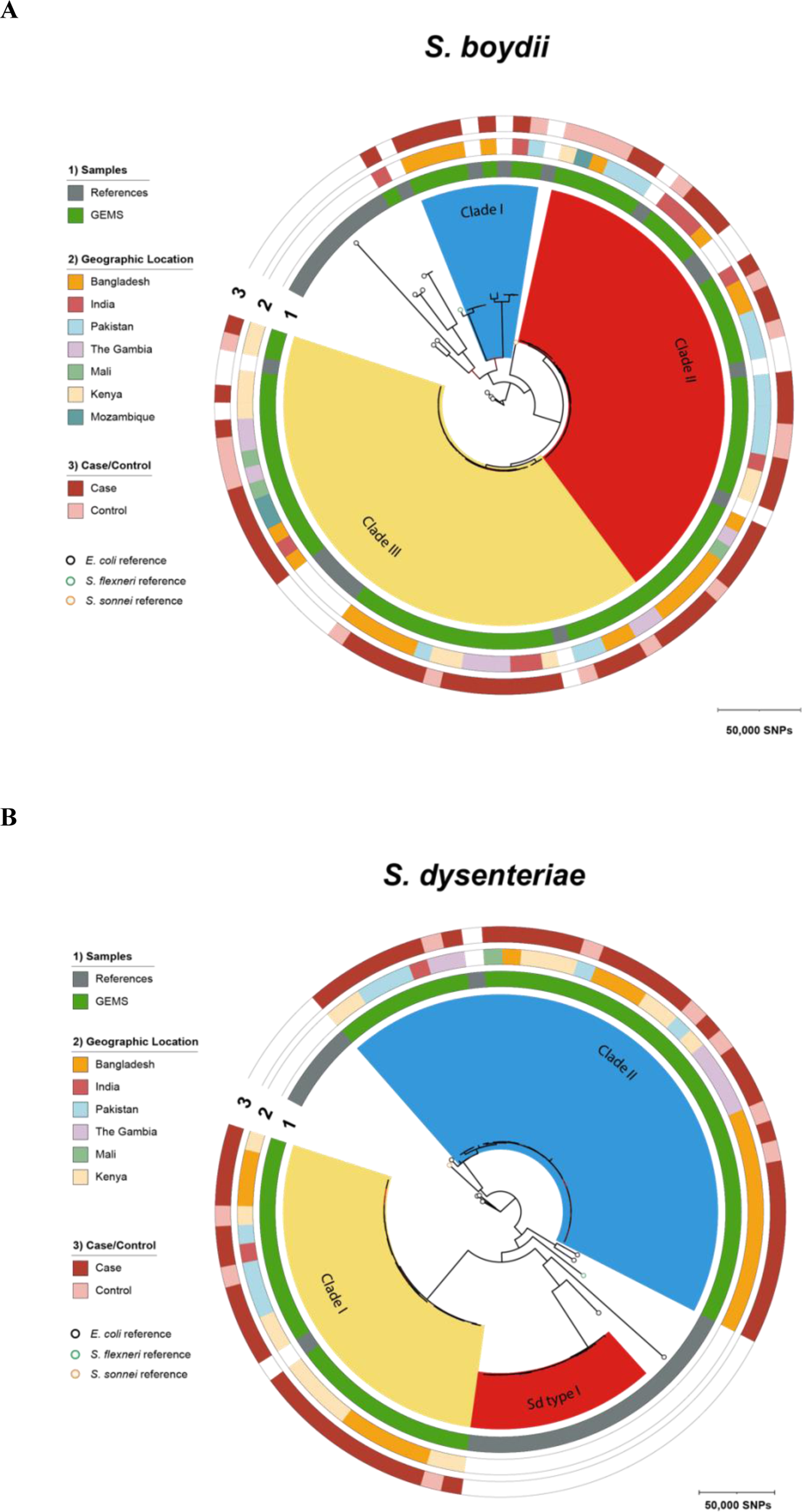
Phylogeny of *S. boydii* and *S. dysenteriae* population from GEMS. ML phylogenetic trees were constructed based on core genome SNPs outside region of recombination from alignments of (A) 79 *S. boydii* and (B) 60 *S. dysenteriae* genomes from GEMS and publicly available genomes. Both trees were rooted using *E. coli* genome. Scale bar represent number of SNPs.

**Fig. S5.**
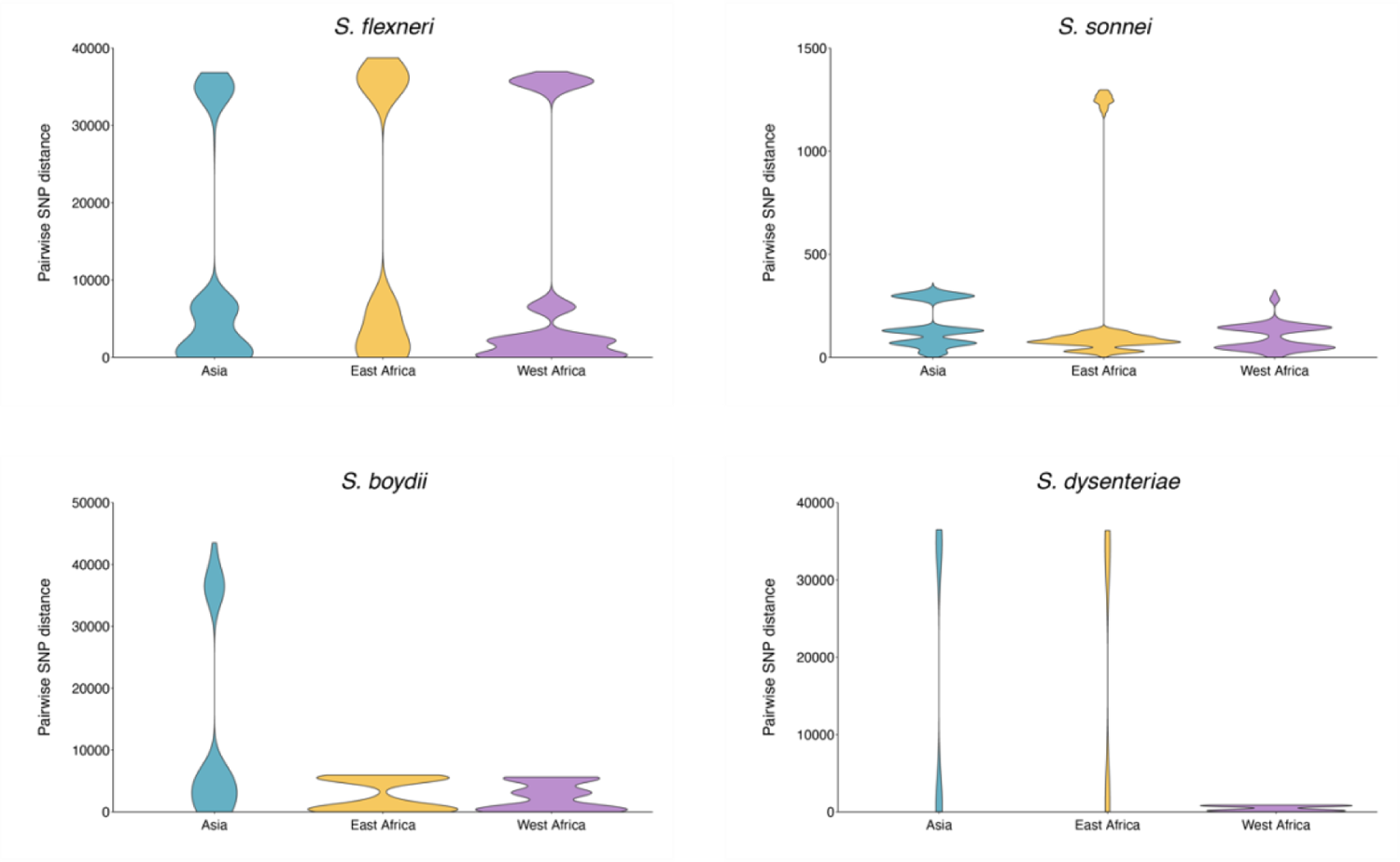
Regional diversity of Shigella spp. Comparison of genomic diversity, as measured by pairwise core SNP distance, across GEMS study sites (Asia: Bangladesh, India and Pakistan; East Africa: Kenya and Mozambique; West Africa: The Gambia and Mali) for *S. flexneri*, *S. sonnei*, *S. boydii* and *S. dysenteriae*.

**Fig. S6.**
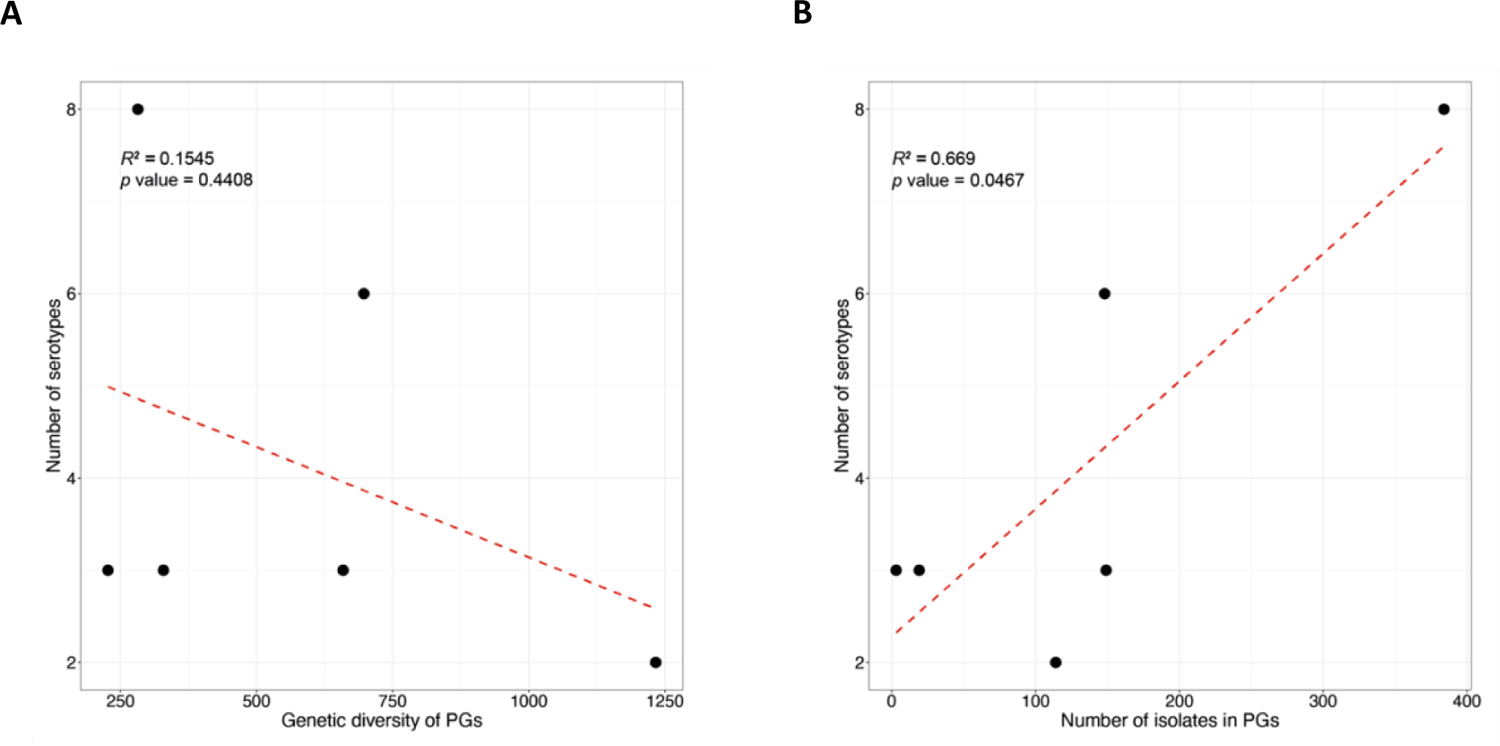
Association of *S. flexneri* serotype diversity with different properties of a genomic subtype. For each of the six subtypes identified among *S. flexneri* (PG1-PG7 and Sf6), the number of different serotypes is displayed along the y-axis and plotted against (A) the number of isolates within the subtype and (B) the genetic diversity of the subtype, as measured by pairwise core SNP distance and plotted along the x-axis. Linear regression analysis was performed to assess the association between serotype diversity and the different properties of subtypes. The regression coefficient of determination (R^2^) and *p*-value are displayed on the top left of each plot.

**Fig. S7.**
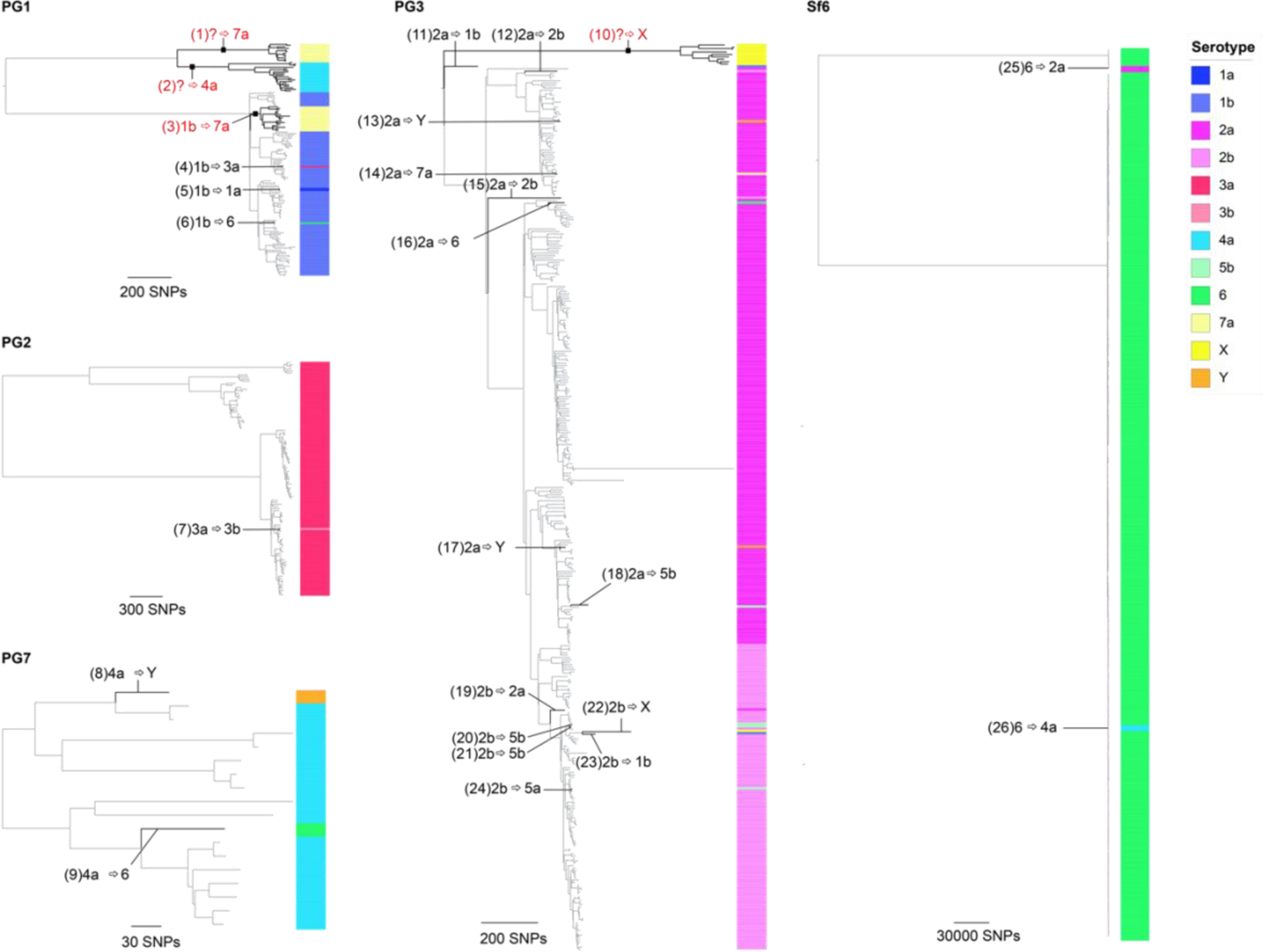
Serotype switching events across *S. flexneri* genomic subtypes. ML phylogenetic tree of each subtype was generated based on core genome SNPs. Serotypes determined through biochemical serotyping are displayed on the right-hand side of each tree, and coloured according to the inlaid key. The 26 inferred serotype switching events occurring along the phylogenetic branches are labelled accordingly. Numbers inside each backets represents switch IDs, with further details provided in table S3. Where the dominant serotype cannot be determined, a question mark is displayed, indicating switch from unknown ancestral type. Serotype switching events resulting in more than two descendant isolates are highlighted in red.

**Fig. S8.**
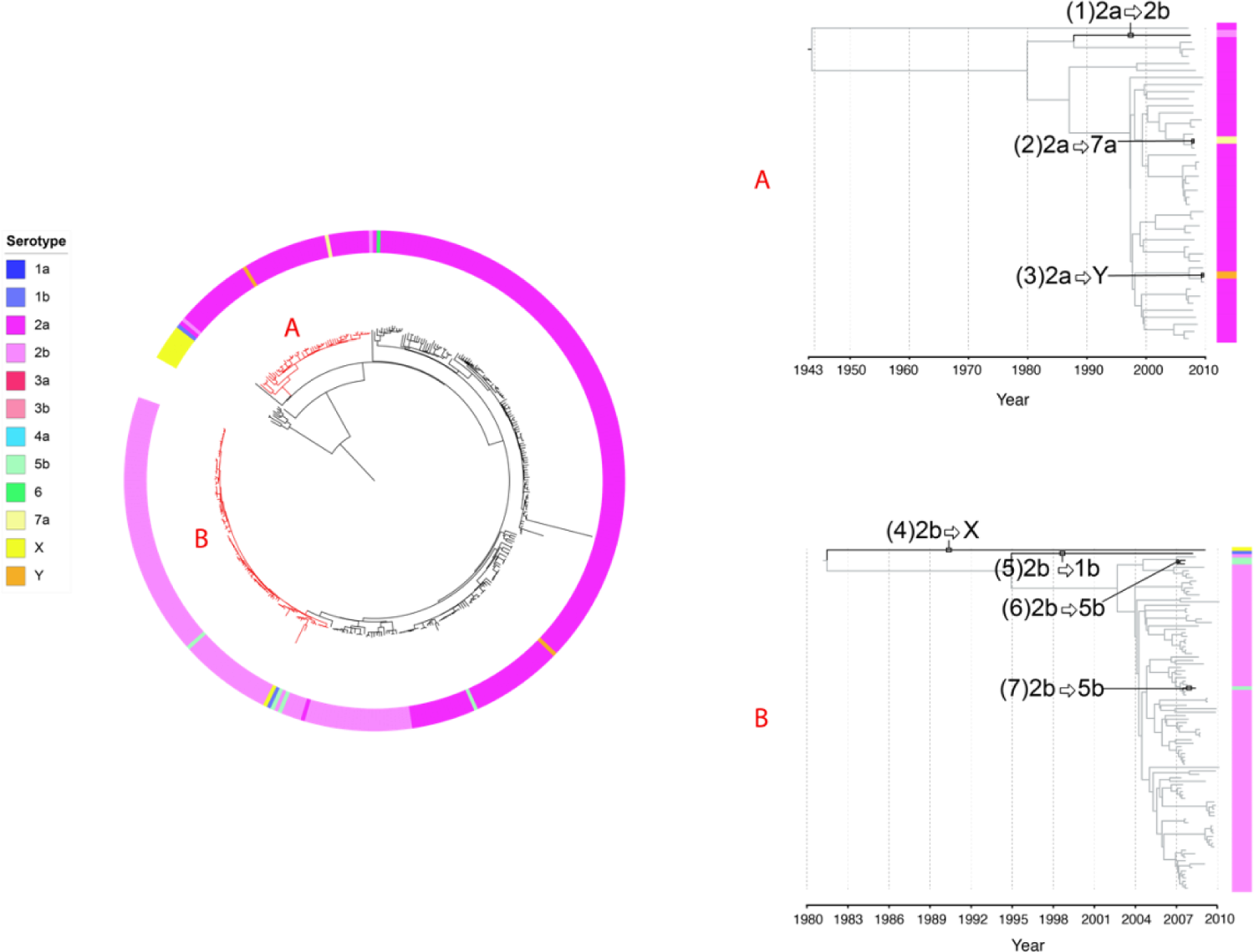
Estimation of time frame for serotype switching among *S. flexneri* PG3 isolates. ML phylogenetic tree of *S. flexneri* PG3 (*n*=384) generated using core genome SNPs is displayed on the right. Isolate serotype is displayed on the outer ring, coloured according to the inlaid key displayed next to the tree. Two subclades with branches highlighted in red were selected for BEAST analysis. Maximum clade credibility trees based on two subclades within PG3 are displayed on the left. Independent switching events occurring along the various phylogenetic branches are highlighted in black, labelled and annotated. BEAST estimated time frame of divergence along the branches of the seven isolates that have undergone serotype switching are shown in table S5.

**Fig. S9.**
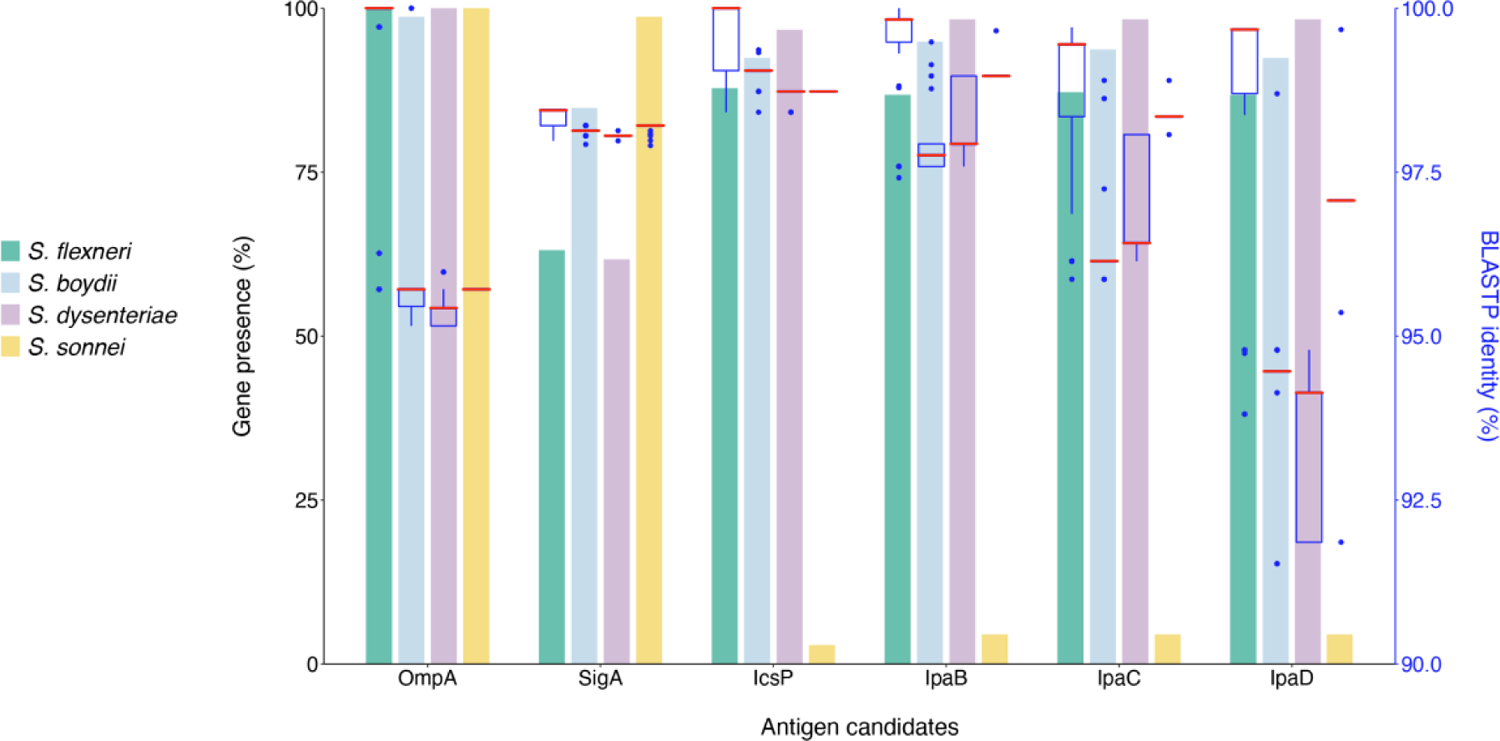
The distribution of vaccine antigen candidate and protein sequence identity among *Shigella* spp. (A) Lefthand y**-**axis refers to the grouped bar plot displaying presence of vaccine candidate genes identified among *Shigella* isolates from GEMS. Bars are grouped by genes and coloured according to species. Righthand y-axis (blue) refers to the boxplot displaying the interquartile range, median (red) and minimum/maximum pairwise percentage identity of the amino acid sequences of antigen vaccine candidates among GEMS, compared against the reference sequences. Presence of genes were identified using BLASTn search against draft genome assemblies and amino acid sequence percentage identity were inferred using BLASTp. (B) Mapping coverage of *Shigella* spp. virulence plasmid. Low percentage of virulence plasmid were detected among *S. sonnei* isolates, likely contributed by the fact that *S. sonnei* virulence plasmid is comparatively unstable and often lost during subculturing.

**Fig. S10.**
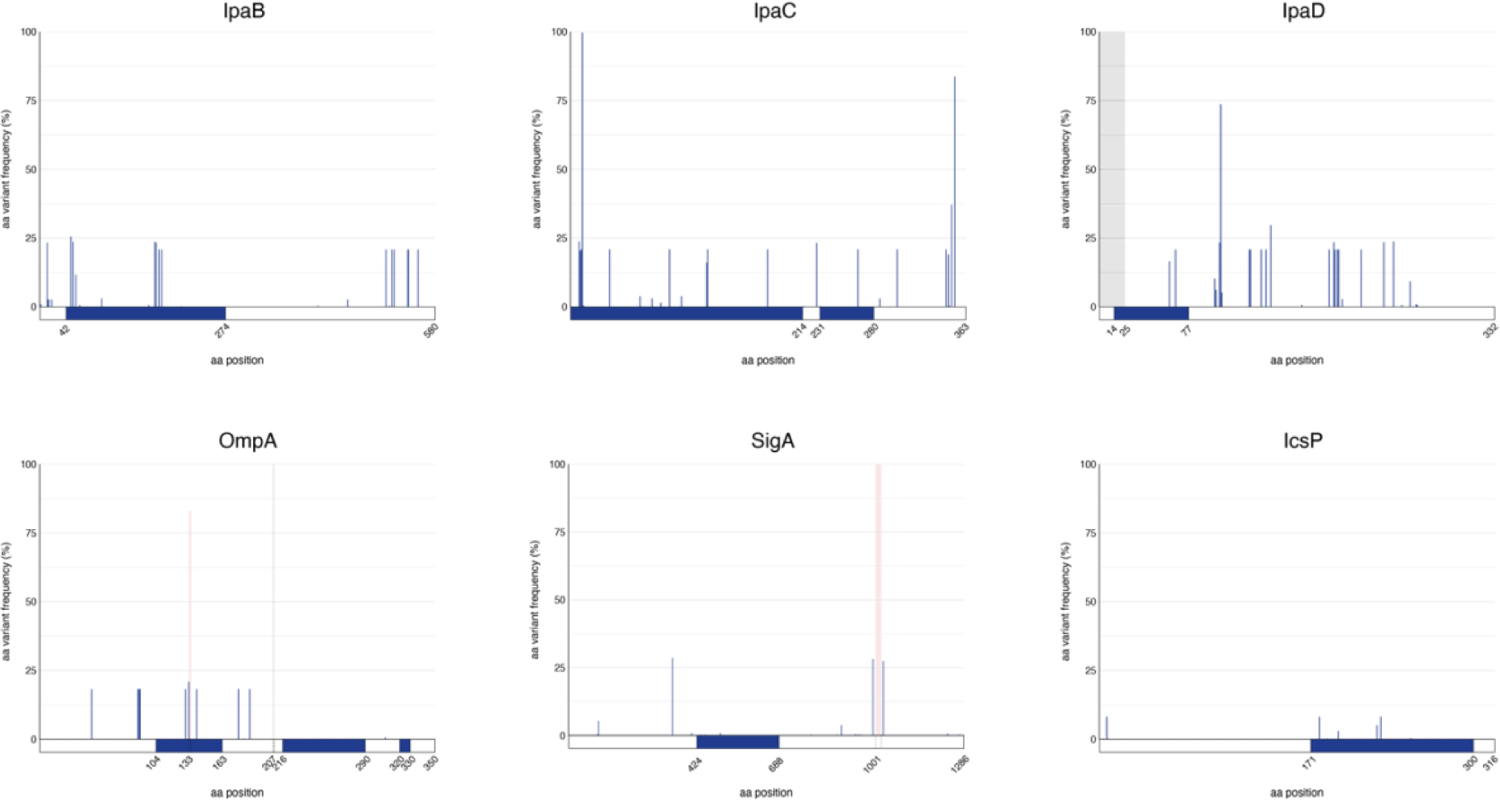
Frequency of amino acid variation among *S. flexneri* population for antigen vaccine candidates. Frequency of amino acid variations within *S. flexneri* genomes for the six vaccine candidate protein sequences. For each protein sequence, the proportion of genomes with the variant is shown along the y-axis with the position of the variant plotted along the x-axis. Grey bars highlight regions where there is a deletion and red bars highlight insertions. Schematic of the known epitope positions (in blue) for the protein sequences are displayed below the x-axis.

**Fig. S11.**
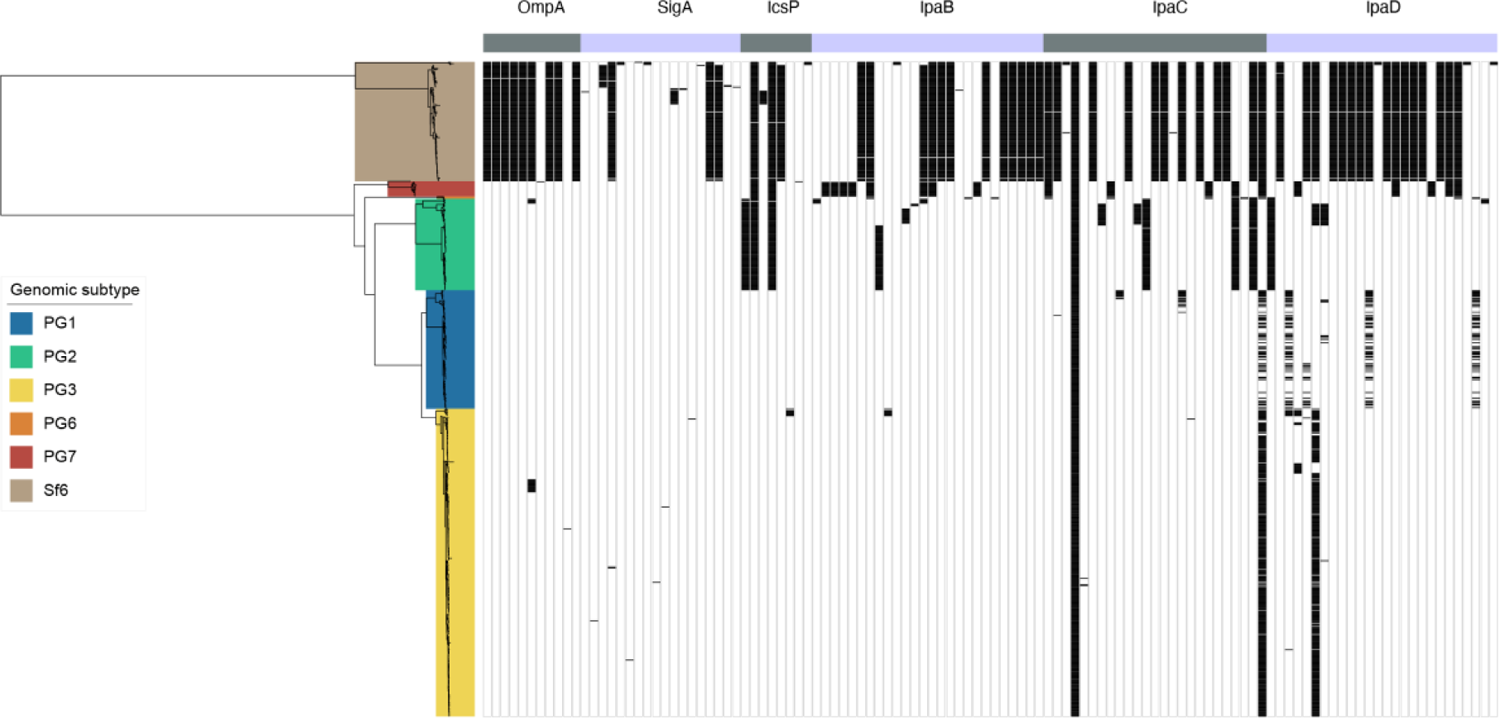
Vaccine antigen variation among *S. flexneri* subtypes. ML phylogenetic tree of 806 *S. flexneri* isolates based on core genome SNPs is displayed on the left, the six subtypes identified among the population are highlighted in different colours according to the inlaid key. The alternating grey and purple colour blocks displayed above the top panel represents the six antigen vaccine candidates assessed in the current study. The matrix in the centre demonstrates presence (in black) of aa variation for each antigen vaccine. Only variable sites are displayed.

**Fig. S12.**
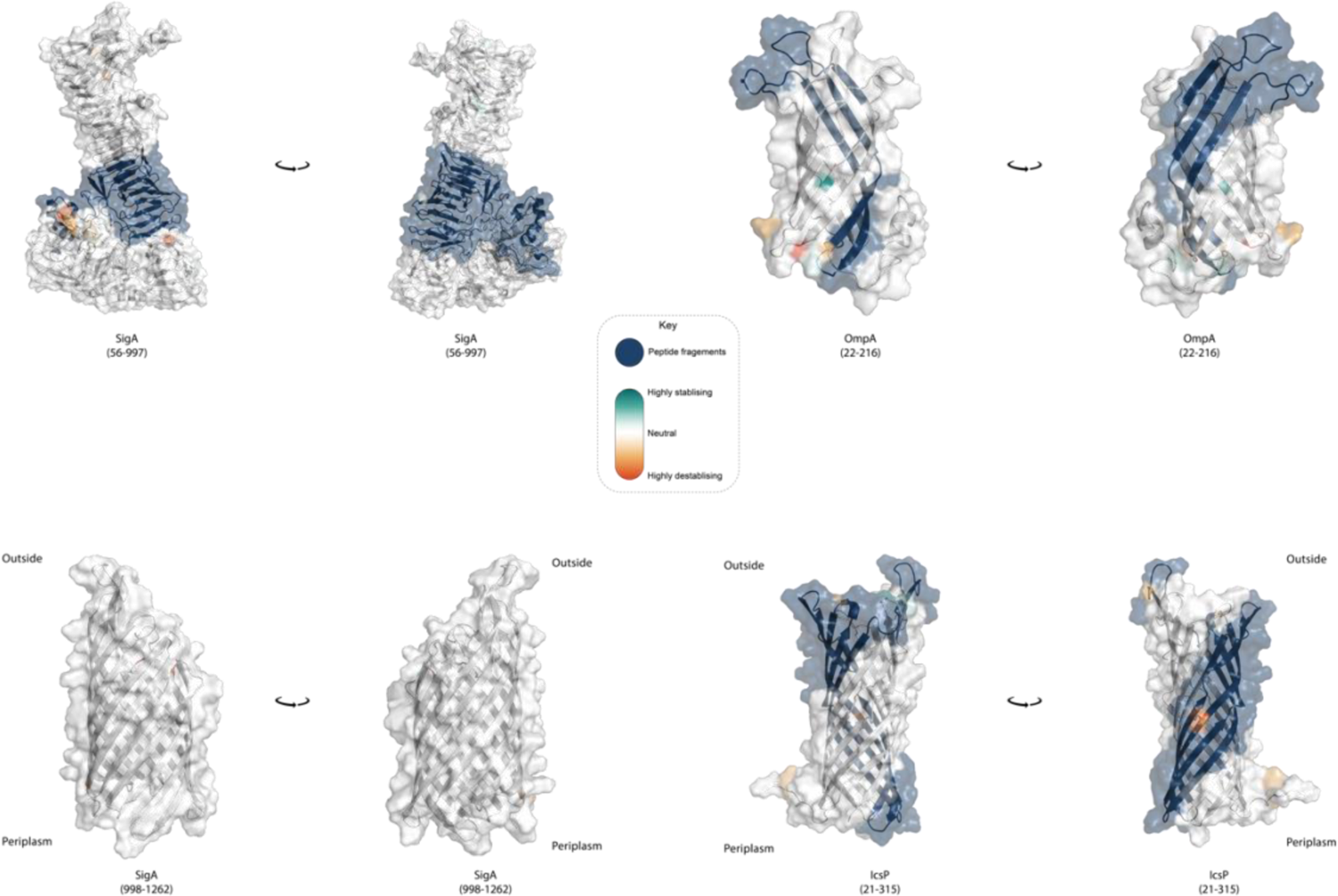
Visualization of mutations on modelled SigA, OmpA and IcsP protein antigens. Visualisation of mutations on modelled proteins, with protein residues modelled shown in brackets. Peptide fragments for OmpA, SigA and IcsP that are used for vaccine development are coloured in blue. Predicted effects of mutations within the proteins are coloured using the scale shown in the key. OmpA, SigA and IcsP are orientated so that the extracellular space is located at the top of the figure, and the periplasmic space is at the bottom.

**Fig. S13.**
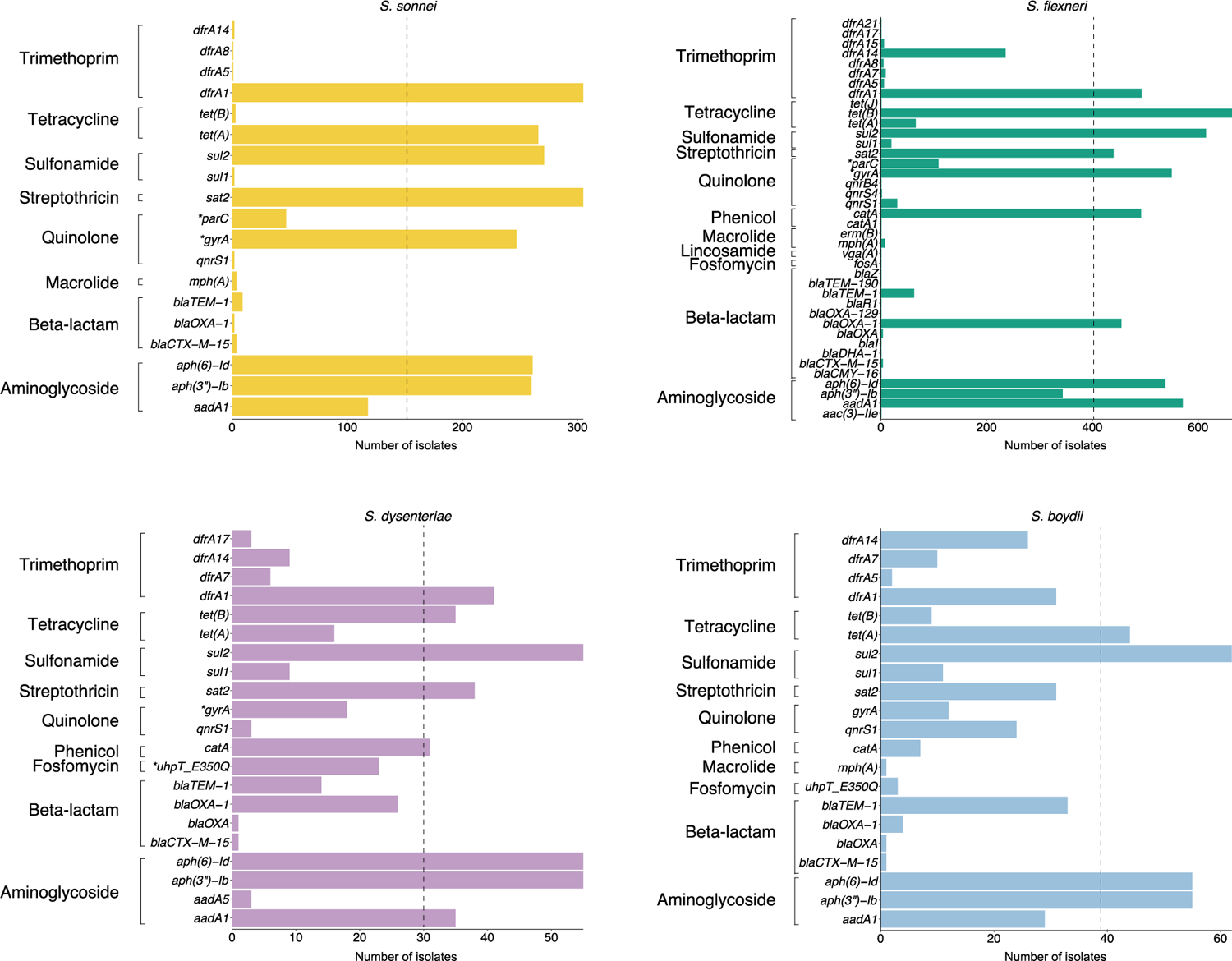
Prevalence of genetic determinants conferring AMR among *Shigella* spp. Bar plots shows the number of genetic determinants detected in *S. sonnei, S. flexneri*, *S. dysenteriae* and *S. boydii* isolates that confer resistance or reduced susceptibility to various antimicrobials. Genes and point mutations (indicated with an asterisk) are plotted along the y-axis and grouped by drug class (displayed on the left). The dashed lines highlight genetic determinants identified in half or more of the isolates for each species.

**Fig. S14.**
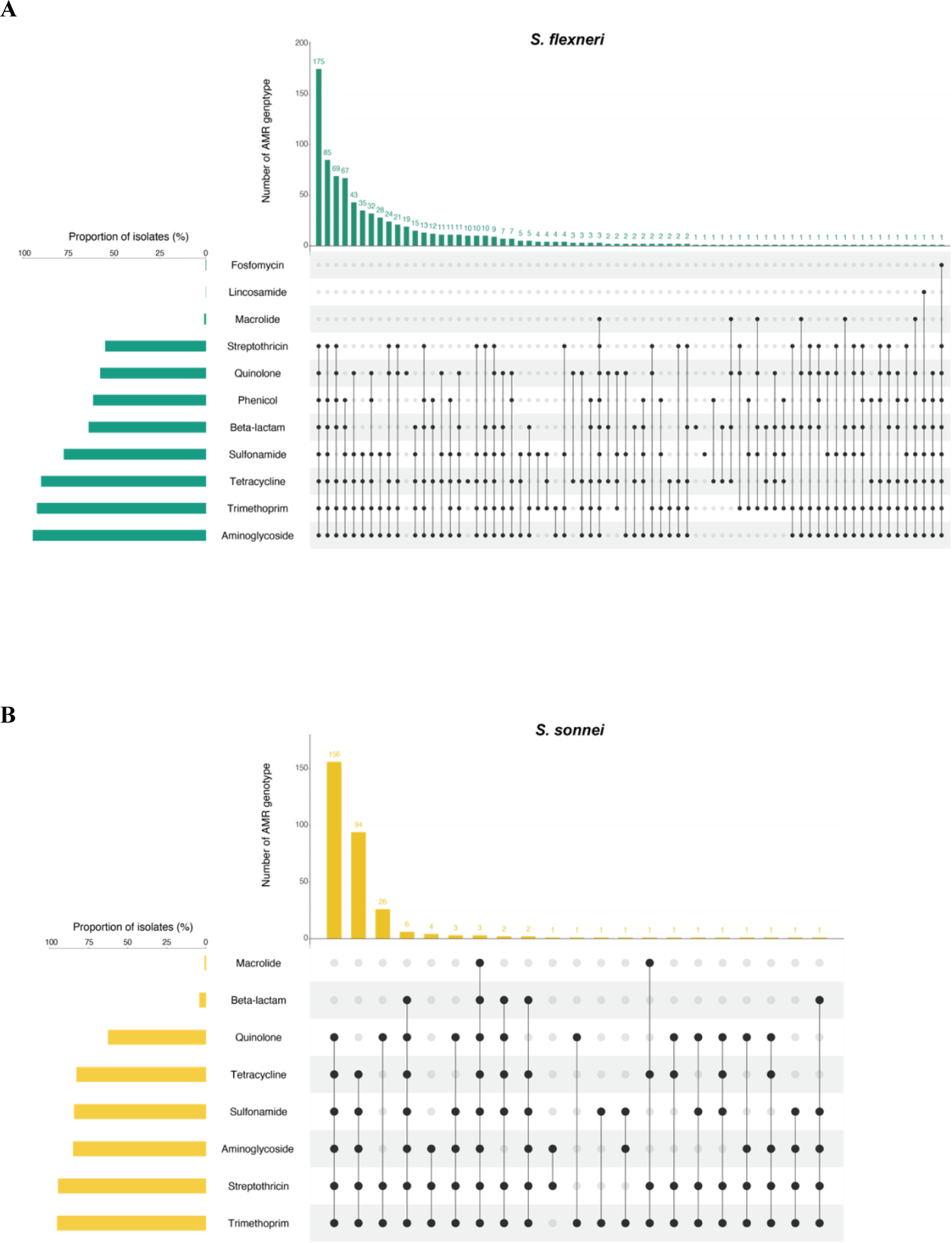
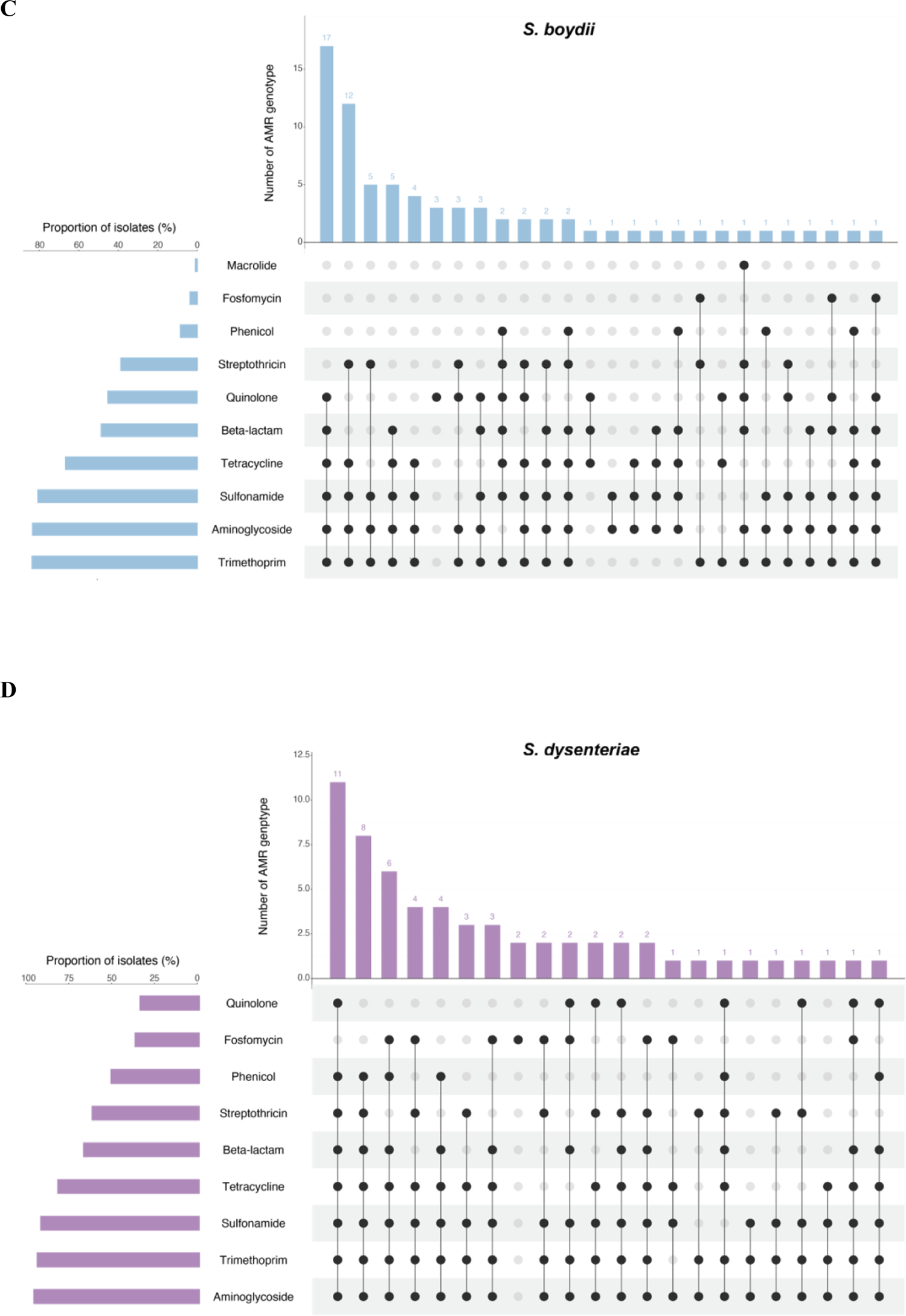
Diversity of AMR genotype resistance profiles. UpSet plots illustrate the AMR genotype resistance profiles for (A) *S. flexneri,* (B) *S. sonnei*, (C) *S. boydii* and (D) *S. dysenteriae.* Genotypic AMR profiles are shown in the combination matrix in the center panel. Each column represents a unique genotypic profile, where each black dot represents presence of a genetic determinant conferring resistance or reduced susceptibility to a drug class (displayed on the left). The vertical the bar plot above the matrix displays the number of isolates with a particular profile, with the exact number of isolates displayed above each bar. The horizontal bar plot on the left of the matrix illustrates the proportion of isolates containing AMR genetic determinants associated with a drug class.

**Fig. S15.**
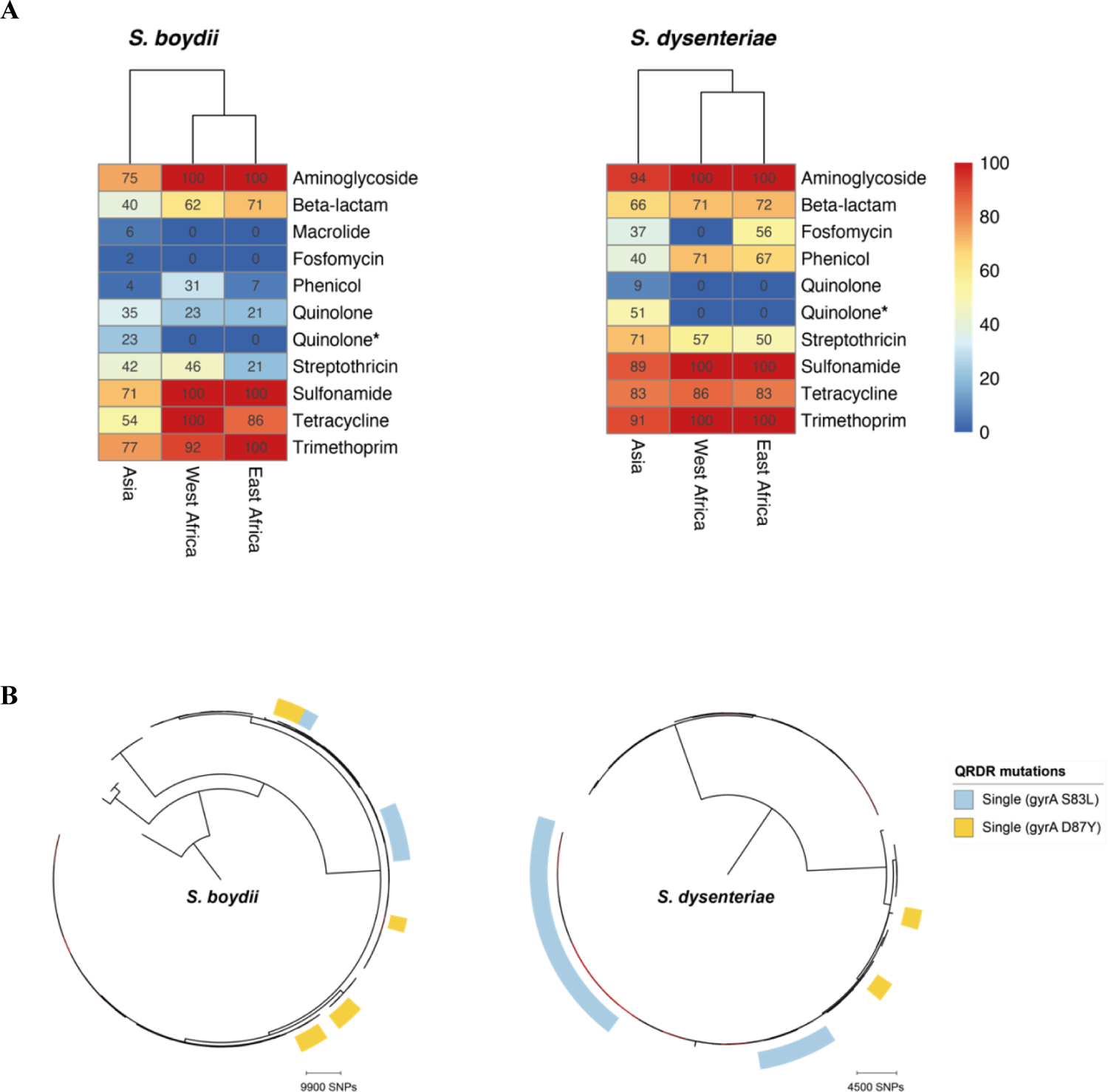
Detection of known AMR genetic determinants conferring resistance (reduced susceptibility marked with asterisk) to various drug class, grouped by region (A) and convergent evolution of ciprofloxacin resistance (B) for *S. boydii* and *S. dysenteriae*.

**Fig. S16.**
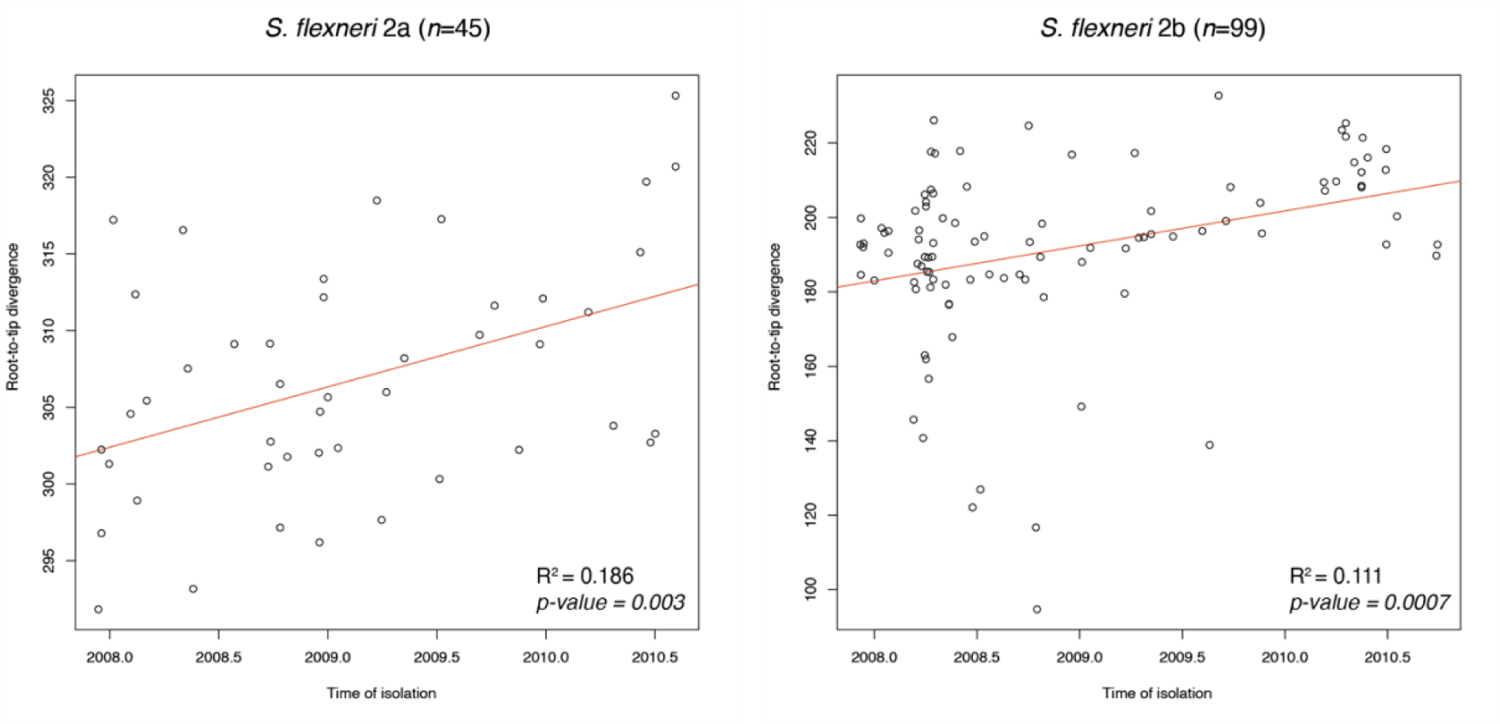
Temporal phylogenetic signal for *S. flexneri*. Correlation between isolate sampling time in months (x-axis) and phylogenetic root-to-tip divergence (y-axis), as estimated by TempEst based on ML phylogeny of each subclade. The two datasets correspond to *S. flexneri* 2a isolates belonging to node A (left) and *S. flexneri* 2b isolates belonging to node B (right) from PG3 in fig. S8. The linear regression line is coloured in red, with the coefficient of determination (R^2^) and *p*-value displayed for each plot.

**Table S1.** Details of Shigella isolates used in this study. Includes accession numbers of the sequencing reads used in the study, *Shigella* serotype, assembly statistics, year and country of isolation, condition of the child (case/control) from which the isolate was derived from as defined by GEMS, genomic subtype, AMR genes and QRDR mutations.

**Table S2.**
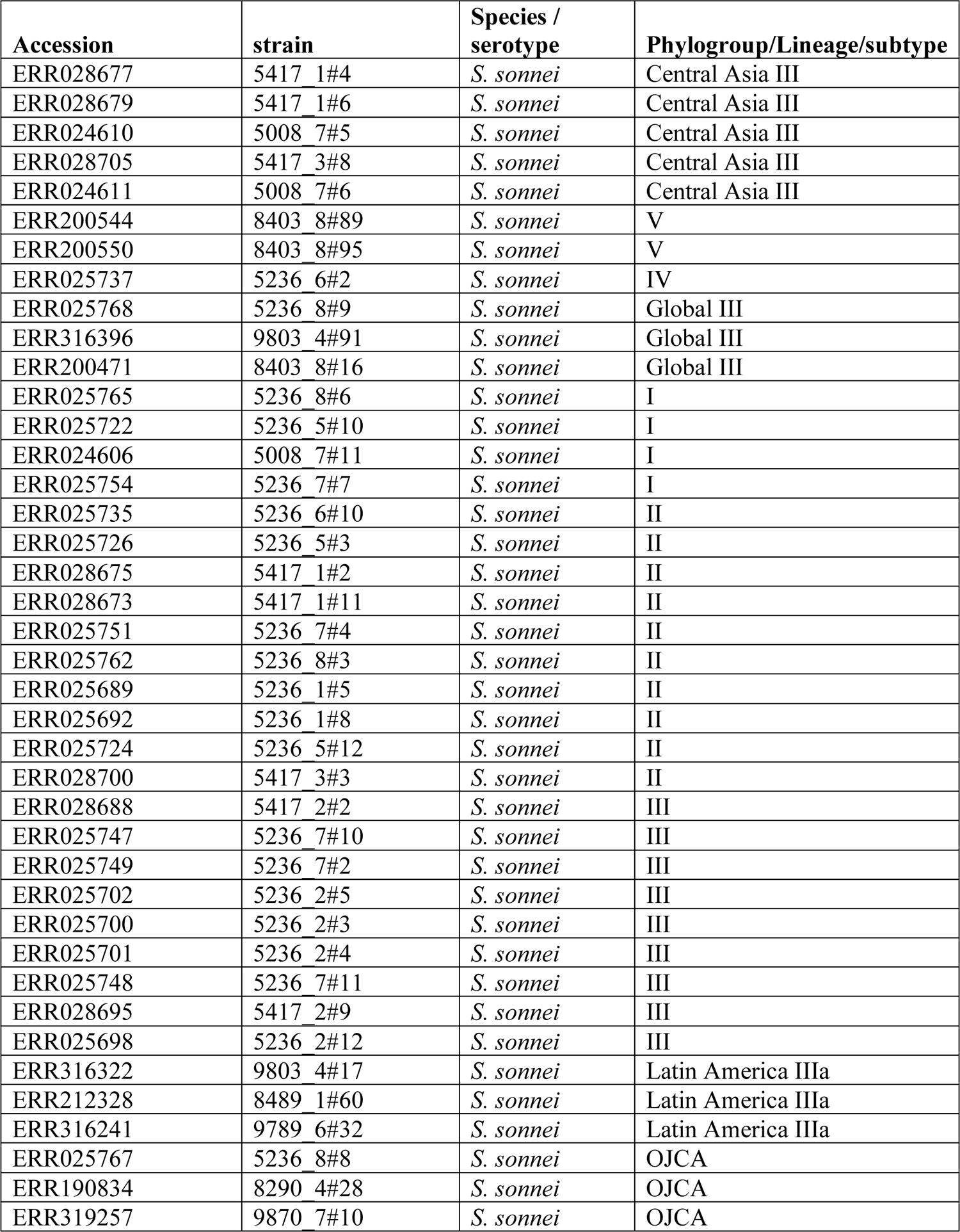
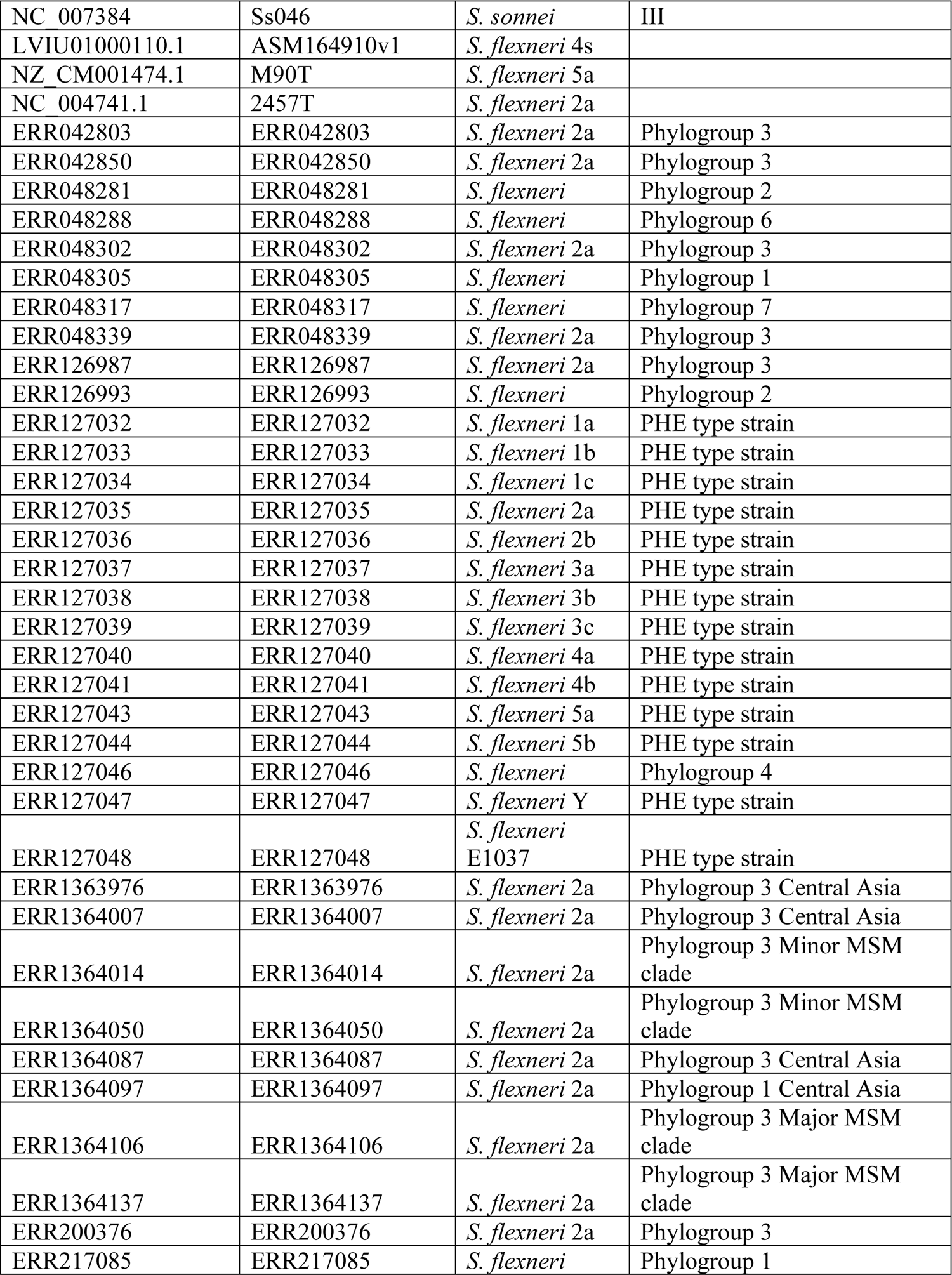
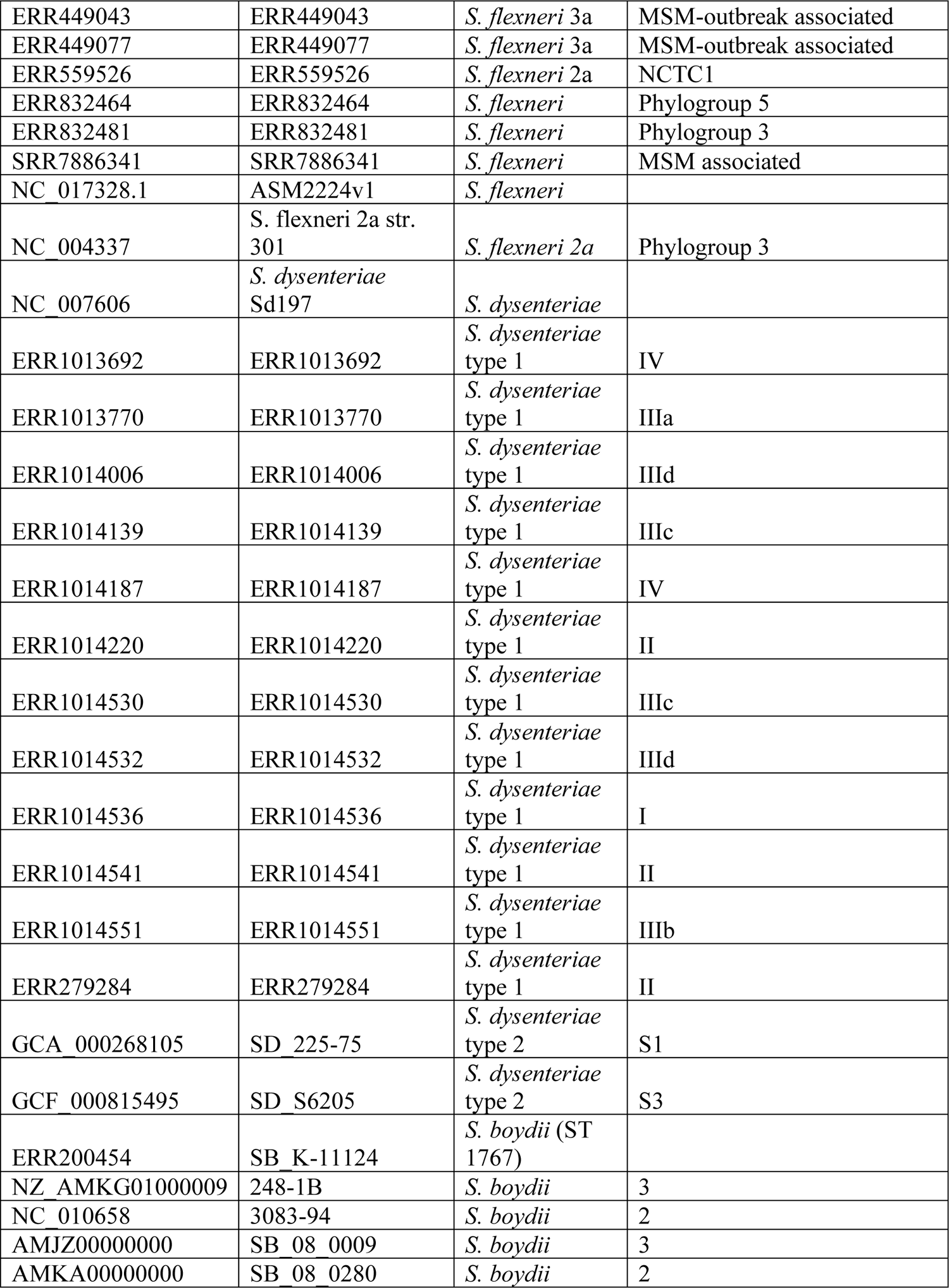
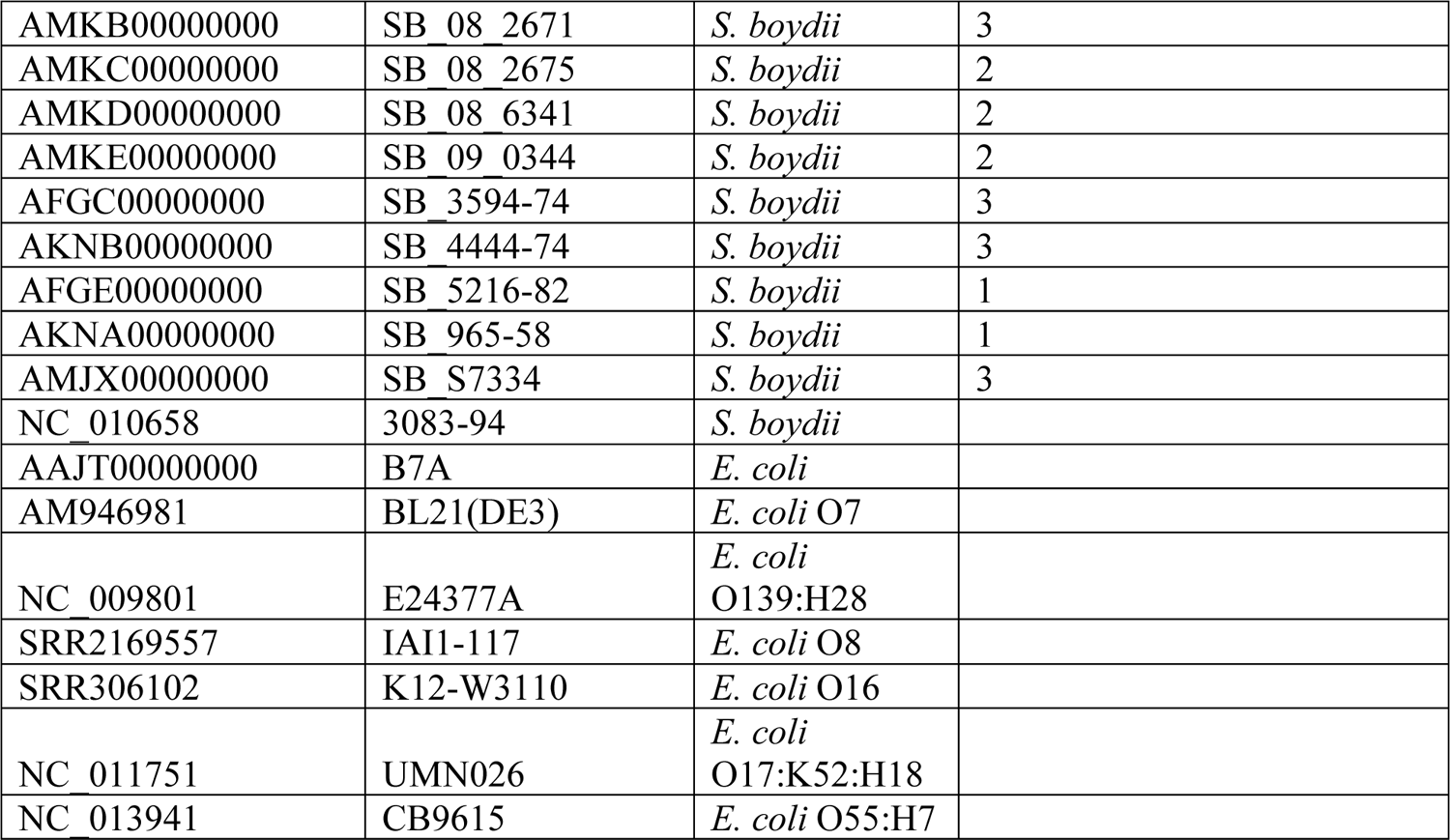
Details of publicly available *E.coli*/*Shigella* genomes used in this study.

**Table S3.**
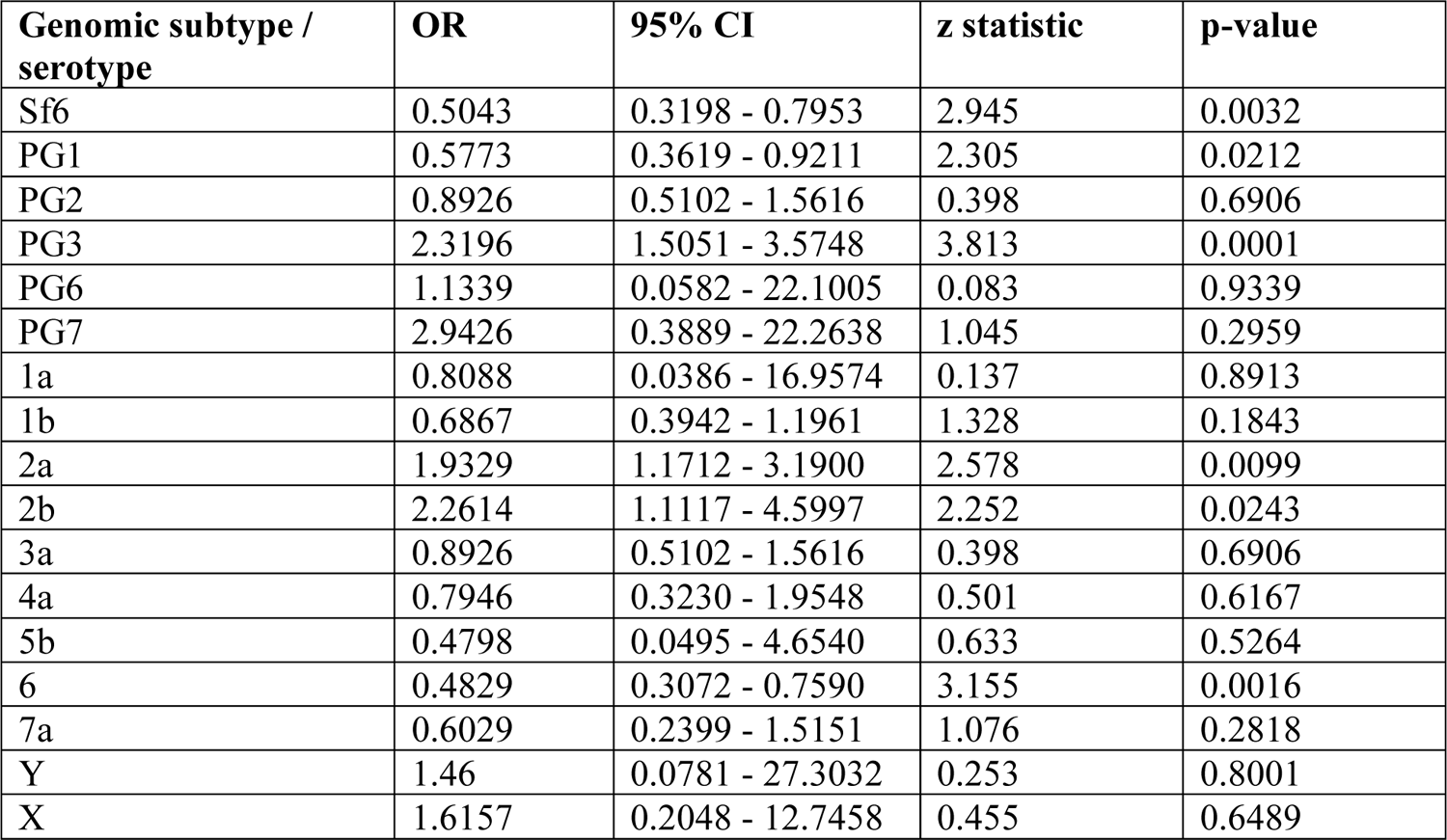
Association of S. flexneri genomic subtype / serotype with case status.

| **Genomic subtype / serotype** | **OR** | **95% CI** | **z statistic** | **p-value** |
| --- | --- | --- | --- | --- |
| Sf6 | 0.5043 | 0.3198 - 0.7953 | 2.945 | 0.0032 |
| PG1 | 0.5773 | 0.3619 - 0.9211 | 2.305 | 0.0212 |
| PG2 | 0.8926 | 0.5102 - 1.5616 | 0.398 | 0.6906 |
| PG3 | 2.3196 | 1.5051 - 3.5748 | 3.813 | 0.0001 |
| PG6 | 1.1339 | 0.0582 - 22.1005 | 0.083 | 0.9339 |
| PG7 | 2.9426 | 0.3889 - 22.2638 | 1.045 | 0.2959 |
| 1a | 0.8088 | 0.0386 - 16.9574 | 0.137 | 0.8913 |
| 1b | 0.6867 | 0.3942 - 1.1961 | 1.328 | 0.1843 |
| 2a | 1.9329 | 1.1712 - 3.1900 | 2.578 | 0.0099 |
| 2b | 2.2614 | 1.1117 - 4.5997 | 2.252 | 0.0243 |
| 3a | 0.8926 | 0.5102 - 1.5616 | 0.398 | 0.6906 |
| 4a | 0.7946 | 0.3230 - 1.9548 | 0.501 | 0.6167 |
| 5b | 0.4798 | 0.0495 - 4.6540 | 0.633 | 0.5264 |
| 6 | 0.4829 | 0.3072 - 0.7590 | 3.155 | 0.0016 |
| 7a | 0.6029 | 0.2399 - 1.5151 | 1.076 | 0.2818 |
| Y | 1.46 | 0.0781 - 27.3032 | 0.253 | 0.8001 |
| X | 1.6157 | 0.2048 - 12.7458 | 0.455 | 0.6489 |

**Table S4.** Details of serotype determining genes facilitating *S. flexneri* (*n*=72) serotype switching.

**Table S5.** BEAST estimated timeframe for serotype switching among S. flexneri PG3 isolates.

| Switch ID<sup>#</sup> | Subclade<sup>¶</sup> | Serotype change | Molecular serotype gene detected<sup>&</sup> | Median branch length (days)<sup>\$</sup> | 95% HPD branch length (days) |
| --- | --- | --- | --- | --- | --- |
| 3 | A | 2a → Y | - | 159 | 16 - 344 |
| 2 | A | 2a → 7a | *gtrII* | 154 | 27 - 307 |
| 1 | A | 2a → 2b | *gtrII*, *gtrX* | 7203 | 4792 - 10009 |
| 7 | B | 2b → 5b | *gtrII*, *gtrX* | 348 | 244 - 479 |
| 6 | B | 2b → 5b | *gtrII*, *gtrX* | 254 | 134 - 491 |
| 5 | B | 2b → 1b | *gtrI*, *gtrII*,<br>*Oac1b* | 4888 | 2962 - 7114 |
| 4 | B | 2b → X | *gtrX*, *gtrII* | 10206 | 5494 - 15408 |

<sup>#</sup> Serotype switching event labelled according to Fig S8
<sup>¶</sup> Phylogenetic subclade isolate belong to
<sup>&</sup> Presence of serotype determining genes, as detected by ShigaTyper, - indicates no genes were detected.
<sup>\$</sup> Phylogenetic branch length represents divergence time, predicted by BEAST and inferred from a time-scaled tree.

**Table S6.** **An overview of the protein modelling**. Table includes information about the antigen candidates modelled, the range of residues the proteins were modelled over, homologues used in template modelling and the QMEAN method and score.

| Species | Antigen candidates | Phylogroup | Serotype | Start | Finish | Homolog | Sequence Identity | QMEAN method | Average local QMEAN score |
| --- | --- | --- | --- | --- | --- | --- | --- | --- | --- |
| *S. flexneri* | OmpA | PG3 | 5A | 22 | 216 | 1QJP | 93% | QMEANDisCo | 0.46 |
| *S. flexneri* | OmpA | PG3 | 5A | 349 | 490 | 1R1M | 41% | QMEANDisCo | 0.31 |
| *S. flexneri* | SigA | PG3 | 2A | 56 | 997 | 3SZE | 44% | QMEANDisCo | 0.71 |
| *S. flexneri* | SigA | PG3 | 2A | 998 | 1262 | 2QOM | 85% | QMEANDisCo | 0.83 |
| *S. flexneri* | IcsP | PG3 | 5A | 21 | 315 | 1I78 | 60% | QMEANDisCo | 0.81 |
| *S. flexneri* | IpaB | PG3 | 5A | 1 | 222 | 3U0C | 76% | QMEANBrane | 0.83 |
| *S. flexneri* | IpaB | PG3 | 5A | 223 | 553 | 3WXX | 19% | QMEANBrane | 0.79 |
| *S. flexneri* | IpaC | PG3 | 5A | 1 | 363 | - | - | QMEANBrane | 0.85 |
| *S. flexneri* | IpaD | PG3 | 5A | 1 | 332 | 3R9V | 100% | QMEANBrane | 0.80 |

**Table S7.** Details of amino acid variants identified for the six antigen candidates among *S. flexneri* isolates from GEMS. Table includes variant type, variant location, reference and alternative variant, and energy score of the variant as predicted by premPS.

